# Primary and recurrent glioma patient-derived orthotopic xenografts (PDOX) represent relevant patient avatars for precision medicine

**DOI:** 10.1101/2020.04.24.057802

**Authors:** Anna Golebiewska, Ann-Christin Hau, Anaïs Oudin, Daniel Stieber, Yahaya A. Yabo, Virginie Baus, Vanessa Barthelemy, Eliane Klein, Sébastien Bougnaud, Olivier Keunen, May Wantz, Alessandro Michelucci, Virginie Neirinckx, Arnaud Muller, Tony Kaoma, Petr V. Nazarov, Francisco Azuaje, Alfonso De Falco, Ben Flies, Lorraine Richart, Suresh Poovathingal, Thais Arns, Kamil Grzyb, Andreas Mock, Christel Herold-Mende, Anne Steino, Dennis Brown, Patrick May, Hrvoje Miletic, Tathiane M. Malta, Houtan Noushmehr, Yong-Jun Kwon, Winnie Jahn, Barbara Klink, Georgette Tanner, Lucy F. Stead, Michel Mittelbronn, Alexander Skupin, Frank Hertel, Rolf Bjerkvig, Simone P. Niclou

## Abstract

Patient-derived cancer models are essential tools for studying tumor biology and preclinical interventions. Here, we show that glioma patient-derived orthotopic xenografts (PDOXs) enable long-term propagation of patient tumors and represent clinically relevant patient avatars. We created a large collection of PDOXs from primary and recurrent gliomas with and without mutations in *IDH1*, which retained histopathological, genetic, epigenetic and transcriptomic features of patient tumors with no mouse-specific clonal evolution. Longitudinal PDOX models recapitulate the limited genetic evolution of gliomas observed in patient tumors following treatment. PDOX-derived standardized tumor organoid cultures enabled assessment of drug responses, which were validated in mice. PDOXs showed clinically relevant responses to Temozolomide and to targeted treatments such as EGFR and CDK4/6 inhibitors in (epi)genetically defined groups, according to *MGMT* promoter and *EGFR/CDK* status respectively. Dianhydrogalactitol, a bifunctional alkylating agent, showed promising potential against glioblastoma. Our study underlines the clinical relevance of glioma PDOX models for translational research and personalized treatment studies.

## INTRODUCTION

Candidate therapeutics for personalized treatment in rare tumors are difficult to test in clinical trials because of inter-tumor differences and the limited number of patients representing specific genetic profiles. Adult diffuse gliomas are a particularly heterogeneous group of rare brain tumors, with grade IV glioblastoma (GBM) being the most malignant subtype^1^. Despite surgery, radiotherapy and chemotherapy the median survival of GBM patients is 14 months and recurrence is inevitable. GBM, characterized as *Isocitrate dehydrogenase* wild type (IDHwt), encompasses tumors with varying genetic backgrounds that affect distinct signaling networks^2,3^, classify into several molecular subtypes with differing expression signatures^4,5^, display variable DNA ploidy^6^ and have different DNA methylation status of the *O-6-methylguanine-DNA methyltransferase (MGMT)* gene promoter. The latter has been shown to predict the response to Temozolomide (TMZ)^7^, the standard-of-care chemotherapeutic agent approved for GBM^8^. A separate group of adult diffuse gliomas characterized by activating *IDH1* (IDH1mut) or *IDH2* (IDH2mut) mutations comprise *1p/19q* intact astrocytomas and *1p/19q* co-deleted oligodendrogliomas, with varying grades (II-IV) and survival rates^9^, further displaying *e.g., PDGFRA* and *CDK4* amplification, *CDKN2A/B* deletion, *ATRX, TP53* or *TERT* promoter mutations^10-12^, as well as a glioma CpG Island Methylator Phenotype (G-CIMP)^13,14^. Several studies point towards an evolution of diffuse gliomas upon treatment and recurrence, where IDH1/2mut astrocytomas show most and IDHwt GBMs least changes in relapsed tumors^15-18^. Still, most identified changes appear idiosyncratic and it remains unclear to what extent current standard treatment leads to molecular changes that could affect drug responses for precision medicine. So far, all targeted treatment attempts in gliomas *e.g*., targeting EGFR^19^, have failed in clinical trials and effective treatment strategies are urgently needed.

A major reason for the numerous failures of clinical trials is the large gap between preclinical models and the treatment situation in patients where the existing preclinical models inaccurately represent human disease. Robust brain tumor models, able to reliably predict the sensitivity of novel personalized treatments in molecularly defined group of patients, are an unmet need^20^. For many years, the glioma research community relied on a handful of longterm adherent cell cultures. Such GBM cell lines undergo significant genetic drift, do not recapitulate certain histopathological features of patient tumors and display inadequate treatment outcomes^21-23^. Some of these shortcomings can be avoided by growing cells in defined serum-free conditions, adapted from neural stem/progenitor cultures^24,25^. These, however, still suffer from a loss of clonal heterogeneity and molecular adaptations to culture conditions^26,27^, in particular loss of focal amplifications^28^. Although patient-derived 3D tumor organoids appear as a robust *in vitro* alternative, lack of the adequate microenvironment and restricted biological material limit their use^29^. For several cancer types patient-derived xenografts (PDXs) established subcutaneously in immunodeficient animals brought a noteworthy advance, as they allow for propagation of primary patient tumors in less selective conditions and retain interactions with non-malignant cells^30^. PDXs were shown to be more accurate in predicting treatment responses than common cell lines^31^. Several international initiatives, such as the EurOPDX and PDXNet consortia, now develop and standardize PDXs for preclinical studies^32,33^. However, a recent evaluation of GBM PDXs highlighted drawbacks in retaining chromosomal copy number alterations (CNAs)^34^ and it remains to be seen whether they represent a sustainable model for testing precision medicine regimens. As subcutaneous PDXs do not recapitulate the natural tumor microenvironment (TME), patient-derived orthotopic xenografts (PDOX) implanted directly in the brain may be more adequate for modeling gliomas in their natural milieu, preserving the physical and physiological constraints of the blood-brain-barrier and the cerebrospinal fluid. To test this, we must assess whether PDOXs can recapitulate patient-specific genetic and epigenetic features, transcriptomic programs and intra-tumoral heterogeneity prior and after treatment, making them amenable as patient avatars for preclinical precision medicine.

We have previously reported that short-term culture of GBM tissue fragments allows for derivation of 3D organoids, while preserving tissue structure and intercellular connections^35^. Intracranial implantation of such organoids in the brain of immunodeficient rodents allowed for conservation of tumor DNA ploidy and major histopathological features such as angiogenesis and invasiveness^36-40^. These GBM PDOXs display clinically relevant responses towards anti-angiogenic treatments^41,42^. Here, we provide further systematic evidence that organoid-based glioma PDOXs are reproducible and clinically meaningful models serviceable for preclinical functional studies. We present a cohort of 40 models generated from primary and paired recurrent gliomas with mixed genetic backgrounds including, amongst others, *IDH1* mutation and distinct *EGFR* variants. We show that these PDOXs preserve key histopathological structures of malignant gliomas (grade III/IV), recapitulate tumor-intrinsic genetic and molecular features at the individual patient level and retain intra-tumoral transcriptomic programs and stem-cell associated heterogeneity. This also applies to PDOX from paired recurrent glioma samples. We further show that glioma PDOXs represent adequate patient avatars for testing precision medicine, also in a high throughput manner. Drug testing in 3D organoids allows for screening *in vitro* at reasonable cost with clinically-relevant responses, which can be further validated *in vivo*. Lastly, we highlight the promising therapeutic potential of Dianhydrogalactitol (DAG), a bifunctional alkylating agent, for treatment of GBM. In summary, our PDOX live biobank represents an important resource for accelerating the development of novel treatment strategies for glioma patients.

## RESULTS

### PDOXs can be generated across diverse clinical high-grade glioma specimen

Fresh tumor samples of 241 glioma patients (189 GBM, 52 grade II-III gliomas) were collected at surgery, including from multifocal samples and longitudinal samples of patients undergoing sequential operations (**Figure 1a-b, Table S1**). Organotypic 3D tumor spheroids, here named organoids, of 300-1000μm were obtained by mechanical dissociation of tissue, without enzymatic digestion, followed by aggregation in short-term culture (max 12 days). This step removes necrotic tissue while preserving a heterogeneous 3D tumor structure, including cellcell interactions, non-neoplastic cells of the tumor microenvironment (TME) and extracellular matrix components^43^, and allows for standardized intracranial implantations. Sufficient material was available for cultures from 72% of collected patient samples, of which 79% GBMs and 68% of grade II-III gliomas presented high quality organoids. Common reasons for lack of healthy organoids were necrotic tissue, tissue damage during surgical procedure (ultrasound) or insufficient material.

**Figure 1.**
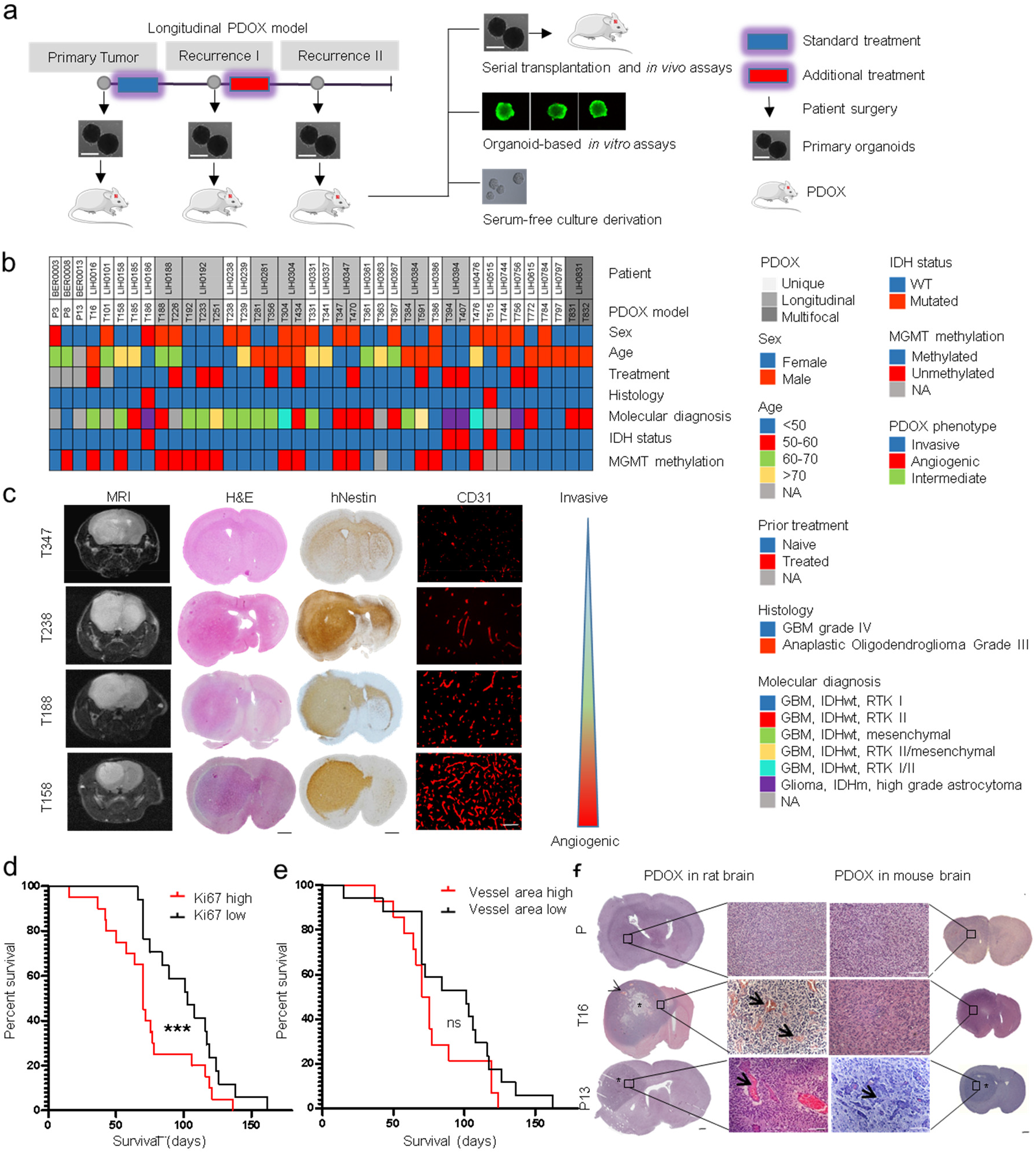
Clinical and histological characterization of glioma PDOX cohort. **a** Schematic of derivation of PDOXs from primary and recurrent patient gliomas. Treatment refers to patients. PDOXs enable tumor expansion via serial transplantation, organoid-based *in vitro* assays including drug screening, genetic manipulations and derivation of long-term *in vitro* cultures. **b** Clinical patient information of corresponding 40 PDOXs (from 32 patients). PDOXs derived from longitudinal or multifocal samples of the same patients are highlighted. See **Table S1** for more information. **c** MRI, Hematoxylin/Eosin, human-specific Nestin and mouse-specific CD31 stainings were performed to assess histopathological characteristics of PDOXs. Representative PDOX models displaying a range of invasive and angiogenic features are shown. Scale bars represent 1mm (black) and 100μm (white). **d** Kaplan-Meier survival curves of PDOXs divided in high and low Ki67 positive cells (mean Ki67 positive cells per model - split by median), ***p_value_ < 0.001 (Wilcoxon signed-rank test). Mean survival of each model ≥ generation 3 was plotted in each group. **e** Kaplan-Meier survival curves of PDOXs divided by vessel area (Average vessel area in μm - split by median value), ns = not significant (Wilcoxon signed-rank test). Mean survival of each model ≥ generation 3 was plotted in each group. **f** Histopathological comparison of the same PDOXs derived in mice or rats. Angiogenic features are amplified in the rat brain (arrows = abnormal vessel morphology, stars = pseudopalisading necrosis, black bar = 1 mm, white bar = 100 μm). Examples are shown for pronounced invasive histopathology (P8), intermediate (T16) and increased angiogenic (P13) growth. Scale bars represent 1mm. See more examples in **Figure S1d**.

Organoids were implanted into the brain of immunodeficient mice (NOD/Scid, NSG) and tumors developed within 4 to 57 weeks depending on the parental tumor (Generation 1). Organoids could also be frozen in DMSO-containing medium for later use. Generally, mice showed longer latency at the first passage (**Figure S1a**) and most PDOXs reached stable tumor development time per patient tumor at Generation 2-4 (**Table S1)**. Successful engraftment and PDOX propagation via serial transplantation (>3 passages) were obtained for 86% of GBMs (35/41, 6 failed due to poor organoid quality), 25% of grade III gliomas (2/8, no association with organoid quality), but none for grade II gliomas. IDHwt GBM organoids survived well a freezing-defrosting procedure, while IDH1mut *(R132H)* gliomas were more fragile, often requiring implantation of freshly prepared organoids or small unprocessed tissue fragments (**Table S1**). Rare activating IDH2mut (*R172K*) gliomas are not yet present in the cohort. Three additional GBMs (PDOXs P3, P8, P13) were initially derived in nude rats^37^ and were further serially transplanted in mice. To date, we have generated a cohort of 40 glioma PDOX models from 32 patients, displaying different clinical characteristics and molecular backgrounds (**Figure 1 b, Table S1**). Our PDOX cohort contains tumors from primary untreated gliomas as well as recurrent tumors after treatment. We obtained paired longitudinal samples, before and after treatment, from 7 patients and were able to generate 15 corresponding PDOXs. One patient (LIH0831) with a multifocal GBM led to 2 PDOX models derived from two tumor tissue collected from distinct locations. Out of 25 PDOX models cultured in serum free medium *in vitro*, 8 glioma stem cell-like (GSC) lines could be propagated long term, including 2 cell lines carrying the IDH mutation (**Table S1**).

### Glioma PDOXs display a range of invasive and angiogenic glioma features

PDOXs derived in immunodeficient mice preserve major histopathological features of patient tumors and display a gradient of invasiveness and vascular pathology depending on the tumor of origin (**Figure 1c**). Angiogenic tumors tended to grow in a more circumscribed manner and showed contrast enhancement on MRI, indicative of blood brain barrier disruption. In line with our previous report^44^, mouse survival was a result of a combination of histopathological features (vascular proliferation, necrosis, invasion) and proliferation index, where high proliferation correlated significantly with poor prognosis (**Figure 1d-e, Table S1**). Models derived from relapsed GBMs showed similar survival and proliferation index compared to treatment-naïve tumors (**Figure S1b-c**). Based on previous experiments with GBM PDOXs generated in rats^37^, we were surprised to find that only a few PDOX models in mice displayed extensive abnormalities in blood vessels. Therefore, we compared identical patient GBMs implanted in either mouse or rat brain. While invasive tumors were similar in mice and in rats, vessel abnormalities were exacerbated in rats in the PDOXs showing only moderate defects in the mouse brain (**Figure 1f, Figure S1d**), including pseudopalisading necrosis, dilated vessels and endothelial cell proliferation. This indicates that the capacity of human GBM to induce angiogenesis is higher in rats compared to mice, likely due to differences in size and cross-species interactions. These interspecies differences were also observed in xenografts derived from serum-free GSC lines (**Figure S1d**).

### Tumor development was independent of non-neoplastic cells present in organoids

Non-neoplastic cells of the TME constituted between 3-25% of all cells in tumor cores in different PDOX models (**Figure S1e**) and these proportions remained stable over serial transplantations. To assess whether the non-tumor compartment present within organoids influenced tumor formation upon implantation *in vivo*, we derived TME-free organoids from FACS-purified tumor cells grown in eGFP-expressing mice and compared them with TME-containing organoids (**Fig. S1f-g**). Both conditions allowed for reformation of 3D organoid structures from sorted cells. Comparison of tumors derived from these two types of organoids showed no significant difference in survival over serial transplantations (**Fig. S1g**). The resulting tumors appeared histologically similar, with the expected level of invasion and presence of an abnormal vasculature. This shows that tumors quickly adapt to the new microenvironment and recreate their niche in the brain by recruiting host-derived TME at each passage.

### Copy number alterations (CNAs) are well preserved in glioma PDOXs

Glial tumors display considerable genetic heterogeneity, with both inter- and intra-tumoral differences^45^. At the level of DNA ploidy, we have previously shown that GBMs present as either mono- or polygenomic tumors and that aneuploidy is a late event in GBM evolution^39^. We found that PDOXs retain patient tumor ploidy states and that both pseudodiploid and aneuploid clones could be propagated by serial implantation (**Table S1**). This is in contrast to long term cultures, where GSC lines of pseudodiploid tumors undergo additional aneuploidization at early passages (**Figure S2a**). Here we show using array-CGH, that at scale CNAs of the parental tumors were maintained with high fidelity in PDOXs both at low and high generations (**Figure 2a, Figure S2b-c, Table S2**). PDOXs clustered next to or in close proximity to their parental tumors. This was also true for longitudinal gliomas, where similar genomic profiles were seen in recurrent tumors after treatment (**Figure 2b, Figure S2d**). Genomic aberrations were also assessed and confirmed by DNA Infinium Methylation EPIC arrays (**Table S2**). Most GBM patients harbored classical genetic hallmarks, such as chromosome 7 gain, chromosome 10 loss and *CDKN2A/B* deletion, which were all retained in PDOXs. This is in contrast to subcutaneous PDXs, where classical GBM CNAs where reported to be lost^34^. Moreover, focal amplicons (*e.g., EGFR, MDM2, MDM4, PDGFRA, MET, CDK4/6*) with the exact same breakpoints were maintained in PDOXs over generations (**Figure 2c, Figure S2c**). IDH1mut gliomas, of which PDOXs could be established, displayed remarkable genomic complexity (**Figure 2d, Table S1-2**).

**Figure 2.**
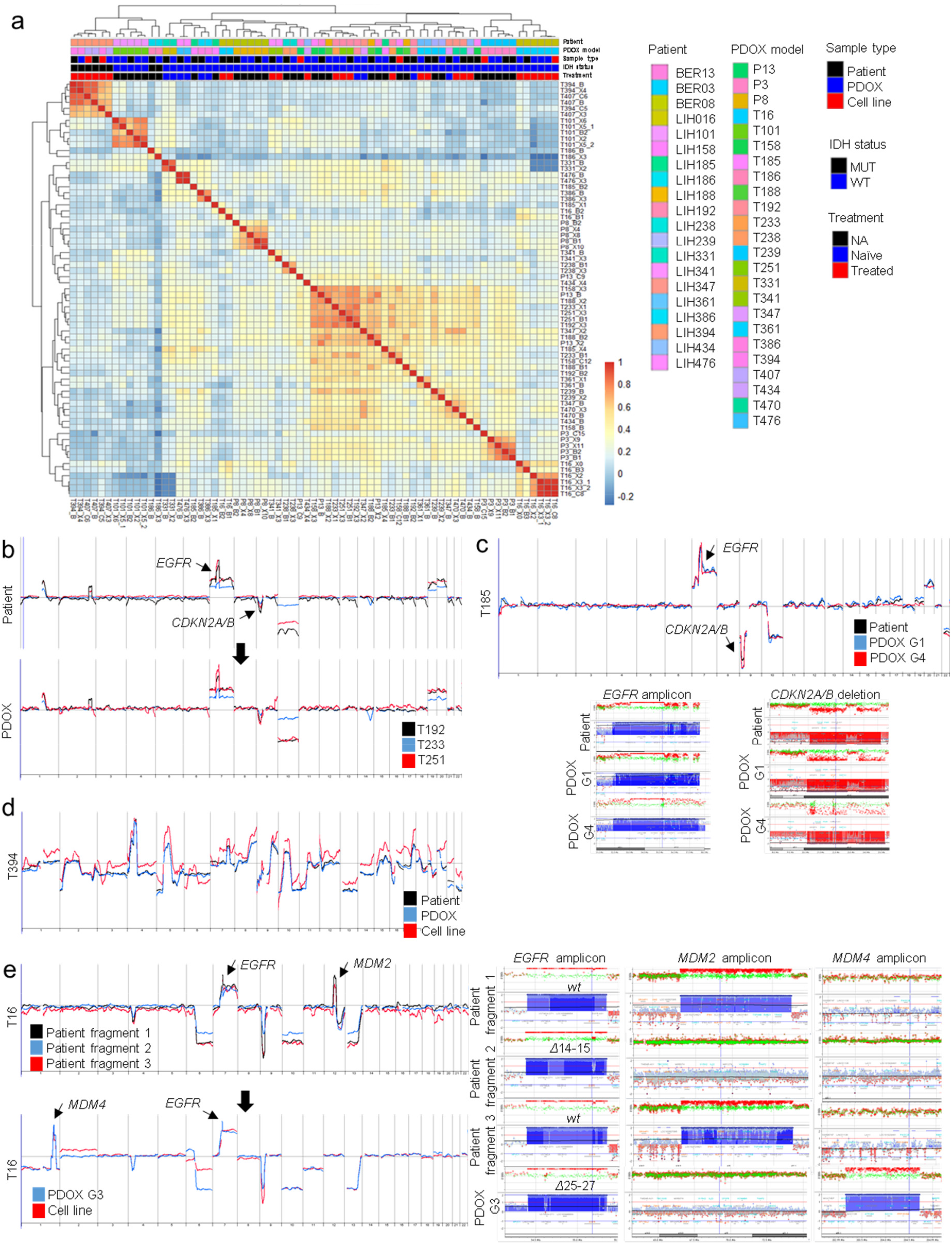
Recapitulation of copy number aberrations in PDOXs. **a** Pearson correlation between patient tumors, PDOXs and cell lines derived based on array-CGH genetic profiles (B = Patient; X = PDOX; C = Cell line; adjacent numbers correspond to passage *in vivo* or *in vitro* respectively). For statistics see **Figure S2b. b** Array-CGH profiles of longitudinal samples (T192-T233-T251) of patient LIH0192 showing retention of genetic aberrations upon recurrence after treatment (radio + chemotherapy). The same profiles were recapitulated in PDOXs. **c** Representative example of an array-CGH profile of GBM patient and corresponding PDOX model (T185 generation 1 and 4). No major changes were detected upon serial xenotransplantation. Identical chromosomal breakpoints are shown for *EGFR* amplicon and *CDKN2A/B* homozygous deletion. See more examples in **Figure S2** and **Table S2**. **d** Example of an IDH1mut glioma patient and corresponding PDOX and cell line showing high genome complexity. Patient was treated with radiotherapy before surgery. **e** Array-CGH profiles of 3 pieces of the same tumor (T16) from patient LIH0016 revealing intra-tumoral genetic heterogeneity (left panels). T16 PDOX and cell line were derived from additional *MDM4/EGFRΔ25-27-amplified* clone. Right panels show the different amplicons in patient tumor fragments and PDOX.

### Rare genetic discrepancies reflect intra-tumor heterogeneity and tumor-specific evolution

It has been suggested that tumors may undergo mouse-specific tumor evolution in subcutaneous PDXs^34^. Here, in our orthotopic xenografts, we only detected minor differences between PDOXs and patients, which could largely be explained by clonal heterogeneity of the parental tumor, particularly at the level of focal amplifications known to be subclonal^46^. *E.g*., patient tumor T16 displayed intra-tumor genetic heterogeneity, where differences in gene amplicons for *EGFR and MDM2* were detected in different tissue fragments dissected from the tumor core (**Figure 2e**). Yet another fragment carrying *MDM4* and *EGFR* amplification with *Δ25-27*structural variant generated the initial 3D organoids and was further propagated *in vivo* over subsequent passages. This was similar for PDOXs T341, P8 and T158 (**Figure S2e-g, Table S2**). These changes may result from tissue sampling bias and selection of specific subclones upon engraftment. Since these were rare, not repetitive and human glioma-specific events, we exclude mouse-specific evolution. Occasionally, we observed acquisition of additional glioma-specific CNAs in later generations (*e.g*., +Chr16 and -Chr 6 in PDOX T101 G6, **Figure S2c**; - 1p21.1-p31.2 in PDOX P13, **Figure S2i**) in line with continuous tumor evolution over time. Loss or acquisition of new aberrations was much more common in cultured GSC lines (**Figure 2e, Figure S2g-i, Table S2**), including loss of *EGFR* gene amplification and protein expression (**Figure S2j**), as noted previously^28^.

### PDOXs recapitulate glioma driver mutations and genetic heterogeneity

To further assess mutation content and clonal architecture of patient tissues and matching PDOXs, we applied targeted DNA sequencing using an extended glioma-specific diagnostic panel (up to 234 genes)^47^. Overall PDOXs showed excellent recapitulation of genetic variants (**Figure 3a-b**). The rare differences between patient tumors and PDOXs were mainly genetic variants detected in patient tumors, but not in PDOXs tissues. These private variants were situated on chromosomes deleted in tumor cells and often had an allele frequency < 50 % (**Table S3)**, suggesting that these variants likely originate from normal human tissue (TME) present in the patients tumor, but not in the PDOX models. Only a handful of genetic variants private to PDOXs were detected and nearly all were located in noncoding regions (**Figure 3a, Table S3**). In comparison, PDOX-derived cell lines showed acquisition of further new variants in cultures (**Figure S3b**).

Targeted sequencing confirmed identified copy-number alterations and further revealed specific mutations characteristic for gliomas **(Figure 3b, Table S4**). Assessed IDH1mut gliomas (PDOX and parental tumor) carried mutations in *ATRX* and *TP53* genes, in line with the molecular diagnosis of astrocytomas obtained by CpG methylation profiling^48^ **(Figure 1b)**. Digital PCR confirmed the presence of wildtype and *R132H* mutated *IDH1* alleles in PDOXs, although variations in ratio were observed probably due to TME signal in patient tumors and tumor aneuploidy (**Figure S3a**). In line with previous reports^27,49^, *in vitro* GSC cultures drastically reduced IDH1 wild-type allele frequency in T394NS, *i.e*. cells had lost wild-type *IDH1* by passage 10, whereas T407NS retained still 20% wild-type *IDH1* allele at passage 13. This was combined with an acquisition of several new variants (**Figure S3b**).

IDHwt GBM PDOXs retained common glioma mutations, including *EGFR, MDM4, PTEN*, *PIK3CA* and *PTCH1* (**Figure 3b**). One PDOX (P13) carried an *IDH2* missense mutation *(W244R)* of unknown significance, not associated with the increased 2HG production (**Table S4**), and thus was considered as IDHwt. *EGFR* gene status was remarkably well preserved. *EGFR* point mutations were detected in the extracellular domain *(A289T, G598V, F254I, R108K)* and co-occurred with *EGFR* amplification. *EGFR* structural variants were present in the extracellular and/or intracellular domains such as *Δ2-7 (EGFRvIII), Δ2-15, Δ6-7, Δ14-15 (EGFRvII)* and *Δ25-27* **(Table S4**). Of note, in agreement with a previous report^50^, PDOX P8 displaying *EGFR A289T* was one of the most invasive and proliferative GBM. In general, our longitudinal models retained similar genetic variants after treatment. Interestingly, LIH0192 patient tumor underwent heterogeneous complex rearrangements leading to a shift from *EGFRvII to EGFRvIII* upon relapse. These changes led to different EGFR protein expression and were retained in the respective PDOXs (**Figure 3c-d**). LIH0347 patient-derived longitudinal PDOXs retained *EGFRΔ*2-15, which also showed immunoreactivity to EGFRvIII antibodies (**Figure S3c**).

**Figure 3.**
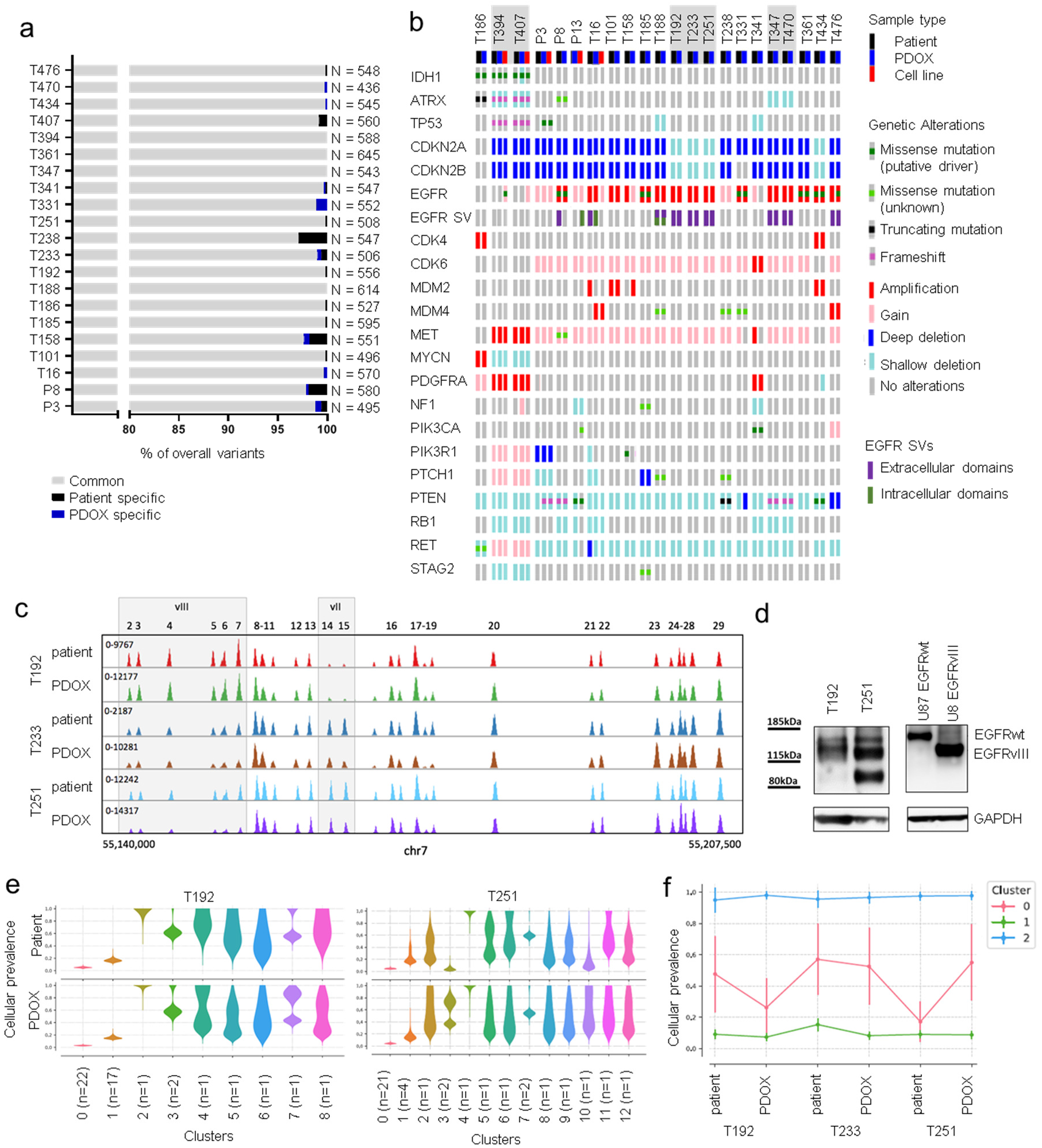
Recapitulation of DNA mutations and structural variants in PDOXs. **a** Recapitulation of overall variants determined by targeted sequencing. PDOXs were compared to respective patient tumors. Number of total variants detected for each patient tumor and PDOXs is displayed. **b** Summary of glioma specific somatic alterations including copy-number changes and mutations in patients and their derivative preclinical models. Samples highlighted in gray represent longitudinal PDOXs. **c** Example of longitudinal GBM samples (T192-T233-T251) of patient LIH0192 showing altered clonal distribution of *EGFR* structural variant *vII* to *vIII* upon relapse, which is recapitulated in the respective PDOXs. Distinct *EGFR* genomic regions deleted in respective variants are depicted. **d** Western blot against EGFR (cocktail antibody recognizing wildtype (wt) and structural variants) confirms protein expression of EGFRwt as well as the respective structural variants *EGFRvII* (in T192) and *vIII* (in T251) with decreased molecular weight. U87 cells overexpressing EGFRwt and *EGFRVIII* are shown for size reference. **e** Cellular prevalence estimates from PyClone representing clonal populations detected in longitudinal patient tumors and respective PDOXs. Examples shown for T192 and T251. Each cluster of mutations was computationally inferred to reflect a subclone. Number of genetic variants contributing to each clone is depicted. **f** Cellular prevalence estimates from PyClone representing clonal subpopulations detected in longitudinal patient LIH0192 and its respective PDOXs. Each line represents a cluster of mutations computationally inferred to reflect a subclone. Only genetic variants detected in all samples were considered for analysis.

We further used PyClone to follow the clonal dynamics upon engraftment of patient tissues and were able to demonstrate that PDOXs retain genetic heterogeneity at the subclonal level (**Figure 3e, Figure S3d**). Subclonal fractions were also retained in longitudinal models of patients LIH0192 and LIH0347, although certain fluctuations in cellular prevalence were observed (**Figure 3f, Figure S3e).** Interestingly, we also observed evolutionary dynamics in *EGFR* amplicons (**Figure S3f)**, arising most probably from evolutionary trajectories of extrachromosomal double minutes^51^. In summary, glioma PDOXs largely recapitulate genetic aberrations and genetic heterogeneity of parental tumors. Rare newly acquired features in PDOXs are glioma specific and may serve as surrogates for the analysis of ongoing genetic evolution in the brain microenvironment, while *in vitro* growth of glioma cell lines leads to additional genetic changes.

### Tumor intrinsic epigenetic profiles are preserved in PDOXs

Cancer-specific DNA methylation patterns are important drivers of gene expression and have been recognized as a preferred prognostic biomarker used for brain tumor subtyping^13,48,52^. Correlation analysis based on EPIC and 450K Illumina DNA methylation arrays showed an overall good correlation between patient tumors and PDOXs (**Figure 4a**), where samples clustered based on IDH status. Although sample type also contributed to the source of variation in the cohort (**Figure S4a)**, IDH status was the main source of variation (**Figure S4a**). IDH1mut gliomas displayed divergent DNA methylation of specific CpG islands compared to IDHwt gliomas (**Figure S4b**). Yet, G-CIMP-low subtype dominated our IDH1mut patient tumors and PDOXs (**Table S5**), presumably in line with their increased aggressiveness^14^. The beta-value distributions were very similar between PDOXs and parental tumors, whereas GSC lines displayed increased DNA methylation at open sea, shelf and shore genomic regions (**Figure 4b**). This was true for IDH1wt and mutated cell lines (**Figure S4c**). Importantly, the *MGMT* promoter DNA methylation status was preserved between PDOX and parental tumor in all but two PDOXs (**Table S5**). Global DNA methylation profiles based on beta-value distributions were also well preserved in longitudinal glioma samples between each other and with their respective PDOXs. Overall, most tumors retained the same DNA methylation profile upon recurrence (**Figure 4c**), including *MGMT* promoter methylation status (**Table S5**), although differences at individual CpG sites are possible.

**Figure 4.**
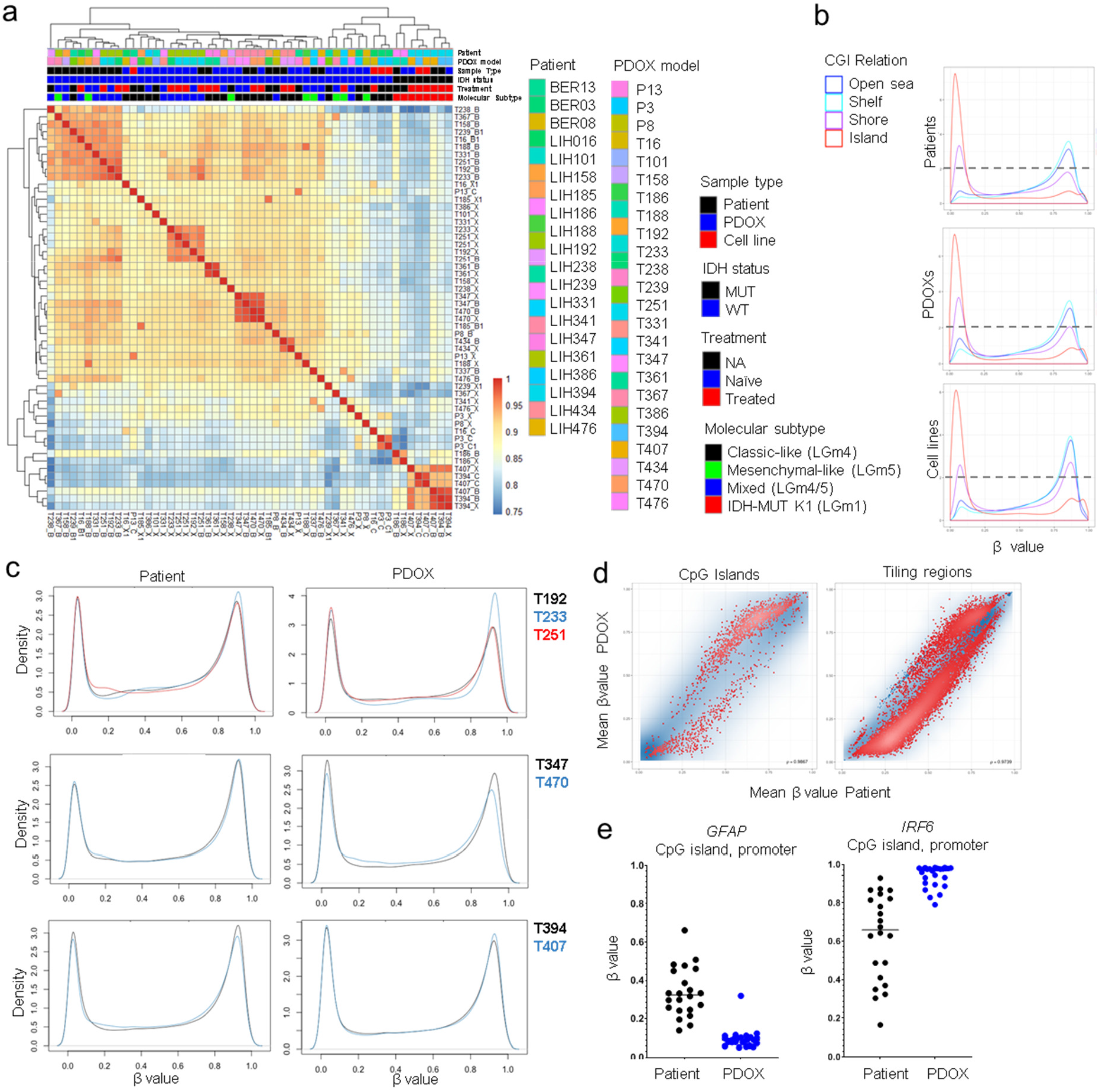
DNA methylation profiling. **a** Pearson correlation of methylation profiles between glioma patient samples, PDOXs and cell lines derived thereof based on 450k and EPIC arrays (B = Patient; X = PDOX; C = Cell line, overlapping regions between arrays only). For statistics see **Figure S4a**. **b** Global beta-value distributions are very similar between patient samples and PDOXs. Cell lines displayed increased DNA methylation at open sea, shelf and shore regions. **c** Beta-value distributions are very similar upon tumor recurrence and are recapitulated in corresponding PDOXs. Examples are shown for longitudinal samples of patients LIH0192, LIH0347 and LIH0394, the latter being IDH1mut. **d** Mean beta-value distribution in patients and PDOXs show increased methylation of a subset of CpG islands and decreased methylation of tiling regions in PDOXs. CpG sites with FDR<0.05 are displayed in red, remaining probes are shown in blue. **e** Examples of hypo- and hypermethylated CpG islands in PDOXs. *GFAP* is widely expressed in GBM, whereas *IRF6* is involved in innate immune response. Differentially methylated sites are changing from hemi-methylated in patient tumors to either unmethylated *(GFAP)* or methylated *(IRF6)* status in PDOXs.

Statistical analysis of paired methylation profiles revealed only minor changes between patient tumors and respective PDOXs (Limma, FDR<0.01). Only 35 individual CpG sites showed differences in mean methylation beta-values above 0.4, corresponding to an essential switch in DNA methylation status, but none were gene annotated CpGs. A partial change of DNA methylation levels (beta-value difference 0.2-0.4) was observed at CpG sites of 226 CpG islands, 89 promoters, 74 gene bodies and 943 tiling regions. Most sites that were demethylated in PDOX corresponded to tiling regions that changed from hemi- to unmethylated (894/943, **Figure 4d**), pointing towards global hypomethylation characteristic for high-grade glioma^52^. This was also true for certain gene promoters specific to GBM cells (*e.g., GFAP*, **Figure 4e**). An increase towards fully methylated CpG sites was observed typically at CpG islands (196/226, **Figure 4d**), including promoters of genes expressed classically by the TME (*e.g., IRF6* for immune cells, **Figure 4e**), reflecting the impact of non-neoplastic cells on methylation profiling^53^. Accordingly, the molecular classification based on previously defined DNA methylation classes^14,48,52^ was well retained in PDOXs (**Figure 4a, Table S5**). Class switches between patient and PDOX occurred from Mesenchymal-like to Classic-like tumors (LGm5 to LGm4, Mesenchymal to RTK II class, **Table S5**), in line with the influence of the TME on DNA methylation profiles as was also shown for gene expression signatures^5^. GSC cell lines displayed more divergent DNA methylation profiles with increased DNA methylation levels (**Figure 4a-b**) and were not clearly classified (**Table S5**).

### Gene expression analysis reveals close resemblance between patient tumors and PDOXs

To determine to what extent gene expression profiles of parental tumors are retained in glioma PDOXs, we performed genome-wide transcript analysis using human-specific microarrays (**Figure 5a**). In parallel, we analyzed cell cultures and corresponding intracranial xenografts from GSC (NCH421k, NCH644) and adherent cell lines (U87, U251). Unsupervised hierarchical clustering revealed close resemblance of PDOXs to corresponding patient tumors, although higher similarity of samples of the same type was observed (**Figure 5a**). Cell lines and their xenografts were more dissimilar and clustered according to their origin, in line with a higher cellular selection and adaptation in long term *in vitro* cultures. Transcriptomic profiles of PDOXs also displayed strongest similarity to GBMs from the TCGA cohort^54^ (**Figure S5a**). Analysis of transcriptional subtypes revealed differences when using the original molecular signatures proposed by Verhaak et al.^4^. However, with the recent tumor-intrinsic classification aimed at reducing the influence of TME^5^, the subtyping remained constant (**Table S6**), suggesting that transcriptomic differences between patient tumors and PDOXs arise from TME-associated gene expression. Cell line subtypes were retained upon *in vivo* growth. Analysis of differentially expressed genes between PDOXs and parental tumors (2-way ANOVA, FDR<0.01, absFC≥2) revealed an increase in tumor intrinsic signals such as cell cycle and DNA repair (**Figure 5b**), which was most prominent in highly proliferative PDOXs (P3, P8, P13, **Figure 5c**). Genes downregulated in PDOXs were associated with TME processes *i.e*., immune response, angiogenesis and macrophage activation (**Figure 5b-c**). Specific markers of human TME components such as endothelial cells (*VWF, KDR*), microglia/macrophages *(ITGAX, AIF1, CD68)*, pericytes/vascular smooth muscle cells (*PDGFRB, ACTA2*) and hematopoietic cells (*CTLA4, CD4, PTPRC*) were depleted in PDOXs (**Figure 5c**). This included also *ABCB1* and *ABCG2*, which we have previously shown to be restricted to brain endothelial cells in human GBM^55^. The general depletion of human TME transcriptome upon xenografting was confirmed by independent component analysis (**Figure 5d**) and flow cytometry (**Figure 5e**). Inter-patient differences were retained in PDOXs, *e.g., EGFR* expression was maintained at similar levels as in patients (**Figure 5c**). We did not detect an upregulation of specific molecular pathways linked to stemness (*i.e*., cancer stem-like profiles), confirming the lack of a particular selection for tumor subpopulations. Indeed, the heterogeneous expression of stem cell markers in GBM, as previously reported^56^, was retained in respective PDOXs (**Figure 5e**) and remained largely stable over serial transplantations (**Figure S5b**). Transcriptomic analysis at the single cell level revealed similar proportions of cycling cells and the presence of a hypoxic gradient in PDOX (**Figure 5f**) as shown for GBM patients^57^. PDOXs also recapitulated intra-tumoral heterogeneity and phenotypic cellular states previously described in GBM patients^5,58^ (**Figure 5g**). Mouse-derived TME, which replaced human TME, showed similar cellular subpopulations as detected in patient tumors including microglia/macrophages, oligodendrocyte progenitor cells (OPCs) and astrocytes comparable to human GBM TME^59^ (**Figure S5c**). In conclusion, our data show that glioma PDOXs recapitulate well tumor-intrinsic transcriptomic profiles. Differences in gene expression signatures at the bulk level can be explained by the replacement of the human TME by mouse cells undergoing GBM-specific adaptation.

**Figure 5.**
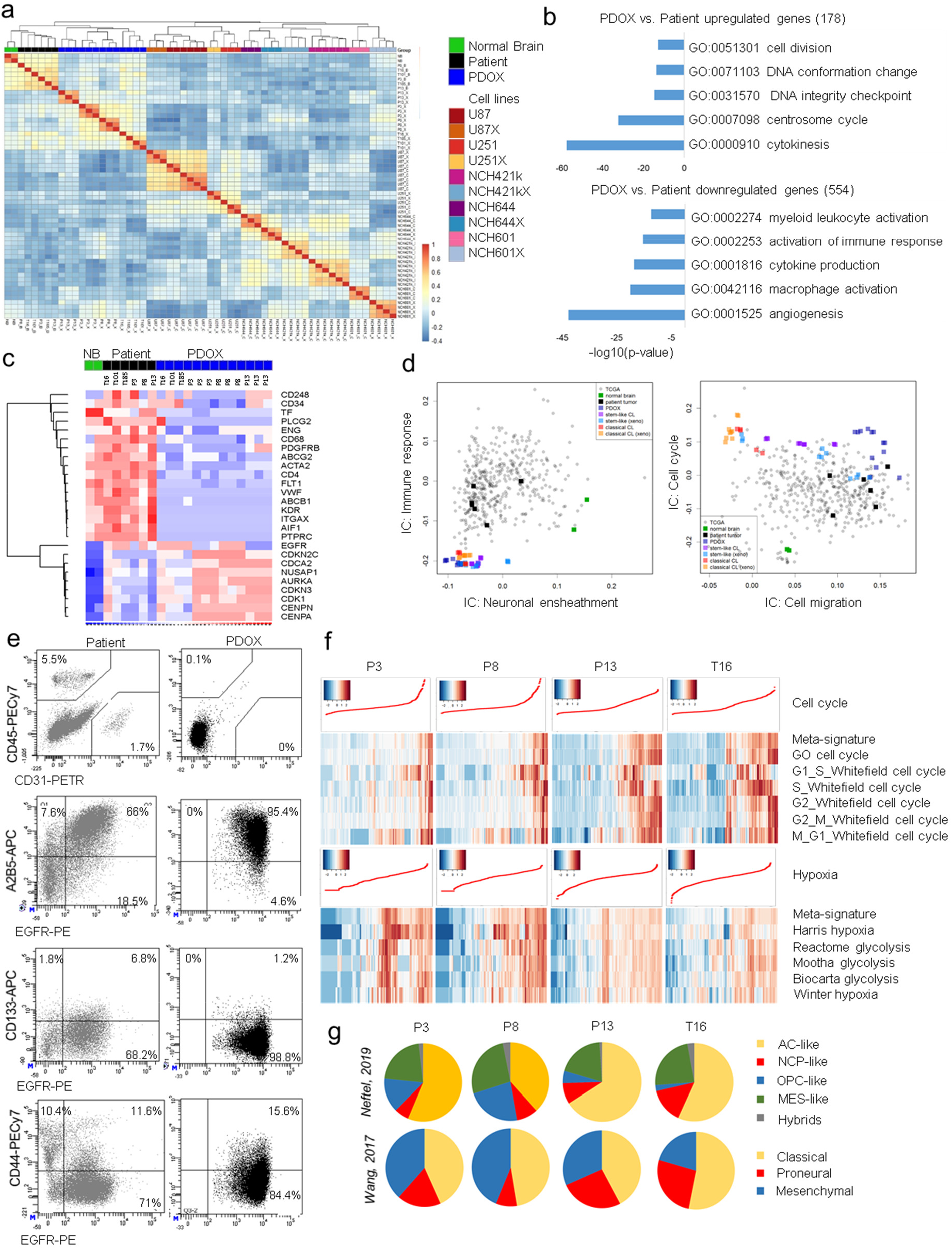
Transcriptomic profiles and intra-tumoral heterogeneity. **a** Pearson’s correlation indicating similarity of genome-wide gene expression profiles between normal human brain, glioma patient samples, PDOXs, GSC lines (NCH421k, NCH644) and classical glioma lines (U87, U251) grown *in vitro* or as xenograft (‘X’). Human specific arrays were applied for transcriptome analysis. **b** Summary of main GO terms characterizing genes differentially present in PDOXs (FRD≤0.01, ab(FC)≥2, Limma). **c** Heatmap representing gene expression levels for a selection of classical TME and cell cycle markers in normal brain (NB), patients and respective PDOXs. **d** Independent component analysis showing depleted transcriptomic signals associated with immune response and neuronal ensheathment in PDOXs and cell lines. Cell cycle independent component (IC) was highest in PDOXs and cell lines, cell migration-associated IC was highest in patients and PDOXs. **e** Flow cytometric analysis to detect human cell subpopulations in patient samples and respective PDOXs. Examples are shown for PDOX T331 expressing EGFR in tumor cells. **f** Single cell signatures showing presence of human tumor cells in distinct phases of cell cycle and hypoxic gradient in PDOXs. **g** Assessment of GBM cellular states^58^ and TCGA GBM subtypes^5^ at single cell level in PDOX tumor cells.

### Preclinical drug testing in PDOX-derived standardized 3D glioma organoids provides clinically relevant outcomes

The PDOX cohort described above constitutes a living biobank maintained by serial transplantation of organoids obtained through mechanical cutting of tumor tissue. This allows to expand the patient tumor in its natural brain microenvironment, generating sufficient material for large scale preclinical drug testing. To this aim we standardized the derivation of uniform 3D GBM organoids amenable for reproducible drug screening *ex vivo*. Organoids were generated from 1000 MACS-purified single tumor cells obtained from PDOXs, which were able to reassemble into 3D organoids within 72h in non-adherent conditions (**Figure 6a**). This allowed for sensitive evaluation of cell viability and toxicity in a 384-well plate format (**Figure S6a**), similar to protocols described for other types of cancer organoids^60^. To assess whether PDOX-derived organoids recapitulate known mechanisms of drug sensitivity and achieve clinically relevant responses, we subjected a cohort of 18 GBM PDOXs to Temozolomide (TMZ), the standard DNA-alkylating agent in clinical practice. Cell responses were calculated as the Area Under the Curve (AUC). In accordance with clinical data, GBM organoids showed only a partial response to TMZ (AUC 200-600, **Figure 6a-b, Figure S6b**). Importantly, tumors with a methylated *MGMT* promoter appeared less resistant in comparison to *MGMT* promoter-unmethylated GBMs (**Figure 6c, S6b**). No differential response was observed between treatment-naïve organoids and organoids derived from patients previously exposed to chemoradiotherapy (**Figure S6c**).

**Figure 6.**
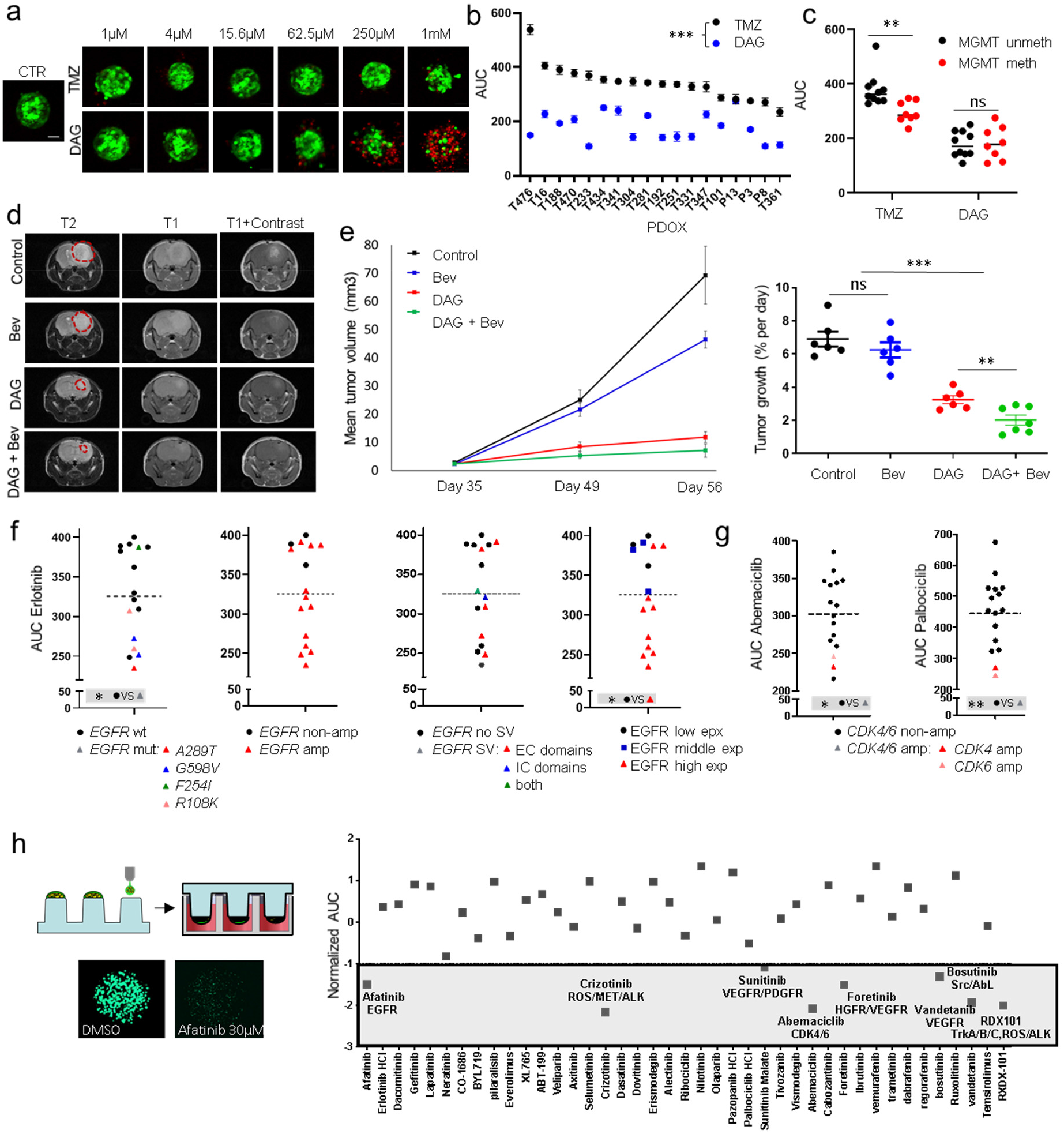
Drug response assessment in glioma PDOXs *ex vivo* and *in vivo*. **a** Drug response was evaluated in PDOX-derived 3D organoids with standardized size (green = viable, red = dead cells). Representative images are shown for TMZ and DAG treatment of T434-derived organoids. Scale bar = 50μm. **b** Quantification of AUC upon exposure to TMZ and DAG. Mean AUC +/- SEM is shown for each model (n = 3). DAG is generally more effective in PDOX-derived organoids in comparison to TMZ (***p_value_ < 0.001, unpaired t-test). **c** Mean AUC upon exposure to TMZ and DAG in *MGMT* promoter methylated and unmethylated tumors. Tumors with methylated *MGMT* promoter show enhanced response to TMZ, while response to DAG is independent of the *MGMT* promoter status (**p_value_ < 0.01, unpaired t-test). **d** PDOX T16 treated *in vivo* with DAG, Bevacizumab or a combination. Tumor progression was assessed by T1 -weighted and T2-weighted MRI images (n = 6-7 mice per group) prior treatment (day 35) and post treatment (day 49 and 56 equivalent of 14 and 21 days since beginning of treatment respectively). **e** Assessment of tumor progression over time reveals significant reduction of tumor growth upon DAG treatment. Tumor growth rate between treatment groups was calculated during entire study (day 35 vs. day 56, n=6-7, ***p_value_ < 0.001, **p_value_ < 0.01, unpaired t-test). **f** Quantification of AUC upon exposure to EGFR inhibitor Erlotinib showing higher sensitivity in *EGFR* mutated tumors. Tumors without *EGFR* amplification and low EGFR expression are most resistant (*p_value_ < 0.05, unpaired t-test); *wt = wild type, mut = mutated, Amp = amplified, SV = structural variant, EC= extracellular domains, IC= intracellular domains, exp = expression*. **g** Quantification of AUC upon exposure to CDK4/6 inhibitors Palbociclib and Abemaciclib shows higher sensitivity of *CDK4* and *CDK6* amplified tumors. (*p_value_ < 0.05, **p_value_ < 0.01, unpaired t-test), *Amp = amplified*, **h** High throughput screening with 42 FDA-approved drug library in PDOX T434. Drug response data is displayed as normalized AUC, ‘-1’ was used as a threshold for strongest hits.

### Dianhydrogalactitol (DAG) exhibits strong efficacy against GBM independent of (epi)genetic background and treatment history

We further tested Dianhydrogalactitol (DAG, VAL-083), a bifunctional compound able to alkylate N^7^-guanine and form interstrand crosslinks and DNA double strand breaks^61^. DAG is known to penetrate the blood-brain barrier and to accumulate in cerebrospinal fluid and brain parenchyma^62^; it is currently tested in clinical trials for recurrent GBM (NCT02717962) as well as in treatment-naïve *MGMT* promoter unmethylated GBM patients (NCT03050736). In our cohort, DAG was significantly more effective than TMZ (**Figure 6a-b)** and the response was not dependent on *MGMT* promoter methylation status (**Figure 6c**). The response was similar in treatment-naïve and relapsed organoids (**Figure S6c**). In view of the strong efficacy of DAG in the *ex vivo* assay we evaluated its ability to decrease tumor growth *in vivo*. Due to the structural similarity with glucose, we hypothesized that uptake of DAG could be further enhanced under hypoxia; we therefore also applied a combination treatment with the anti-angiogenic agent Bevacizumab previously shown to induce hypoxia in GBM^41,42^. As expected, Bevacizumab treatment did not halt tumor progression despite decreased contrast enhancement on MRI (**Figure 6d**) and blood vessel normalization (**Figure S6d**). DAG monotherapy led to a dramatic reduction in tumor growth (**Figure 6e**) as assessed by MRI, which was further reduced by combined treatment. Histological assessment of tumor-containing brains confirmed the strong reduction in tumor volume upon DAG treatment (**Figure S6e**). This was paralleled by an increase in DNA damage in tumor cells, determined by H2AX phosphorylation (H2AX-P) (**Figure S6f**). Limited H2AX-P was also seen in normal brain cells close to the meninges and the subventricular zone, but to a much lower extent than in tumor cells. In summary, we show that DAG has a consistently favorable drug profile against GBM, thus representing a promising candidate for GBM treatment either alone or in combination with anti-angiogenic compounds.

### PDOX-derived organoids are amenable to high throughput drug screening for precision medicine

To evaluate the potential for personalized treatment regimens of our models, we functionally assessed the response against a set of EGFR/ErbB small-molecule tyrosine kinase inhibitors (Erlotinib, Gefitinib, AZD3759, AG490, Daphtenin) and CDK4/6 inhibitors (Abemaciclib, Palbociclib) with varying specificity in 16 PDOX-derived organoids with variable genetic makeup of this pathways. The inability to preserve gene amplification and *EGFR* structural variants in most cell culture models including GSCs^28^, did so far not allow for accurate personalized preclinical studies. Our testing group included GBM with different status of *CDK4, CDK6* and *EGFR* amplification, *EGFR* genetic variants and point mutations (**Figure 3b, Table S4**). The responses against EGFR inhibitors were highly variable across organoids (**Figure 6f, Figure S6g)**. In contrast to kinase domain mutations found in lung cancer, glioma-specific extracellular domain mutants are known to respond poorly to EGFR inhibitors^63^. Still, we found that GBM carrying *EGFR* mutations, except for *EGFR F254I* (PDOX T434), were more sensitive to Erlotinib (**Figure 6f**), but not to other EGFR inhibitors (**Figure S6g).** This is in accordance with the fact that *EGFR R108K, G598V* and *A289T* are missense mutations leading to a gain-of-function, shown previously to sensitize tumors to Erlotinib^64^. The role of *EGFR F254I* is currently unclear. *EGFR* amplification and corresponding high protein expression also had an impact on the sensitivity to Erlotinib, where non-amplified tumors with low protein expression were most resistant. EGFR structural variants did not sensitize tumors in our cohort to any of the five compounds. Similarly, tumors carrying *CDK4* (PDOX T434) and *CDK6* (PDOX T341) amplification were most sensitive to CDK4/6 inhibitors Palbociclib and Abemaciclib (**Figure 6g**).

Finally, we performed a proof-of-concept study on PDOX-derived organoids for high throughput drug screening using the cell printing technology based on ASFA Spotter ST^65^. PDOX T434 derived GBM cells were dispensed onto pillars (1000 cells per pillar), embedded into alginate drops and allowed to reform 3D organoids (**Figure 6h**). A library of 42 FDA-approved drugs was then applied for 7 days and response was assessed via a High Content imaging system (CV 8000) recognizing viable cells. To select the strongest hits, we applied normalized AUCs (Z score, −1 threshold)^63^. The screen showed similar responses as the 384-well plate protocol (**Figure S6h**), and confirmed sensitivity of T434 tumor cells to Abemaciclib and resistance to Erlotinib and Gefitinib. Interestingly, it revealed sensitivity to several other inhibitors, including Afatinib, - a second-generation EGFR inhibitor. In summary, we show that PDOX-derived GBM organoids display clinically relevant drug responses and can be applied for personalized drug screening in a high-throughput manner.

## DISCUSSION

Although major discoveries can be performed directly on patient tumors, biological material is restricted, limiting such studies to descriptive analyses and low-throughput preclinical assays. Here we present a living tumor biobank that encompasses the clinical diversity of high-grade diffuse gliomas. 40 PDOX models were established from treatment-naïve glioma patients and patients that underwent standard-of-care, of which 15 represent paired longitudinal models. Our PDOX cohort contains tumors of varying genetic and molecular background, and represents a valuable tool for drug screening, functional studies and *in vivo* drug efficacy studies. We show that glioma PDOXs recapitulate (1) glioma tissue architecture, including features of angiogenesis and invasion, (2) genetic variants and CNAs, including rare gene amplifications (3) epigenetic and transcriptomic tumor intrinsic signatures, (4) intra-tumoral genetic, transcriptomic and stem-cell associated heterogeneity, (5) clinically relevant drug responses. To our knowledge, this is the first at scale cohort of orthotopic glioma xenografts with comprehensive analysis at the molecular level and proof-of-concept treatment responses that are clinically meaningful. Our models and associated molecular data are openly shared and available at the PDXFinder portal (https://www.pdxfinder.org/) and via the EurOPDX consortium (https://www.europdx.eu/). They represent a robust tool for reliable expansion of patient tumor material while maintaining close identity with the parental tumors, and allow for high-throughput drug testing and precision medicine.

Most glioma PDX models are derived by subcutaneous implantation of tumor fragments^66,67^, which may not accurately mimic the complex and distinctive brain microenvironment. As direct implantation of tissue fragments to rodent brains is challenging, GBM orthotopic xenografts usually rely on single cell dissociation followed by *in vitro* cultures prior to xenotransplantation si^,66,68-70^, where cultures are often maintained for unspecified time and passage number. To minimize the loss of tissue architecture and heterogeneity, which may lead to clonal selection and adaptation^71,72^, we derive organoids from mechanically minced glioma tissue, which is only briefly maintained in culture without any *in vitro* passaging. This process retains the primary nature of GBM cells and as a result it does not allow for indefinite growth of organoids *in vitro* as seen under serum-free conditions^29^. In our hands, most GBMs and lower grade gliomas give rise to short term organoids. Successful PDOX establishment is, however, largely limited to IDHwt GBMs and to grade III and IV IDH1mut gliomas. This is in concordance with the general selection of aggressive tumors upon PDX generation in different tumor types, including pediatric brain tumors^73^. So far, only a handful of IDH1mut glioma models were described, which all suffer from long development time and/or changes in *IDH1* status^67,74–78^. Successful IDH1mut models in our cohort were defined molecularly as high-grade astrocytomas with abundant chromosomal aberrations, *CDKN2A/B* loss, *ATRX* and *TP53* mutations and G-CIMP-low signature. These molecular features correspond to the most aggressive IDH1mut gliomas^14,18,77^. Importantly, our models retain *R132H IDH1* heterozygosity and efficient production of 2HG^79^. *In vitro* cultures derived from these tumors either died or led to depletion of the wildtype *IDH1* allele, in line with previous reports^49,80^, suggesting that IDH1mut gliomas require components of the brain microenvironment to maintain their growth. Importantly, our fully annotated cohort displays a wide variety of genetic features not recapitulated in other models (*e.g., EGFR* and *PDGFRA* amplification), thus reflecting the wide inter-patient heterogeneity of high-grade gliomas. Our PDOX biobank also contains 15 unique paired models derived from the same patients at initial diagnosis and upon disease relapse.

The recapitulation of histopathological features of gliomas has been challenging with classical serum-grown cell lines, as they largely lose the characteristic invasive potential of diffuse gliomas upon xenotransplantation^81,82^. Infiltrative growth is maintained in all our PDOXs, although the extent of typical glioma features, including invasion, angiogenesis and proliferation rate can greatly vary across models, likely reflecting inter-patient heterogeneity. We find that prominent angiogenic features are rare in mice compared to rats, which may arise from differences in brain size and/or in molecular interaction between species, suggesting that for studies targeting angiogenesis, rat PDOX models may be more appropriate. Others also reported gradients of invasive and angiogenic features across GBM xenografts, with limited endothelial proliferation and necrosis in mouse brains^67,77^, while large subcutaneous tumors displayed extensive angiogenesis^67^.

We have previously reported that GBM organoids and corresponding PDOXs models faithfully retain tumor cell ploidy^39^. Here we further show that PDOXs accurately maintain distinct genetic backgrounds of parental tumors, including gene amplifications of *EGFR, PDGFRA, MDM2/4 and CDK4/6*, which are difficult to derive and preserve *in vitro*^27,28^. PDOXs also recapitulate complex *EGFR* variants and mutations present concomitantly with *EGFR* amplification. At scale, we found that individual genomic profiles are highly conserved in PDOXs. We did not detect major divergences in CNAs as reported for subcutaneous GBM PDXs^34^. The difference in results may be related to the subcutaneous transplantation, which may lead to a different tumor evolution than in the brain. Alternatively, it may be due to differences in data analysis, since array-CGH based CNA determination, employed by us, is known to be more accurate than CNAs inferred from gene expression profiles^83^. We have further observed extensive preservation of genetic intra-tumoral heterogeneity, although some fluctuations in subclonal architecture were detected. Interestingly, we report a case of *EGFR* variant selection, observed both upon tumor relapse in patients as well as upon xenografting. This may be linked to high levels of *EGFR* amplification and the presence of extrachromosomal double minute structures, which are known to show evolutionary dynamics^51^.

In rare cases PDOX models showed engraftment or expansion of specific genetic clones, with distinct gene amplifications or mutations, differing from the originating tumor. Genomic events that were private to the PDOX correspond to classical glioma aberrations, known to be heterogeneous late events in GBM^84,85^, supporting the notion that the PDOX-dominating clones were a result of original intra-tumor heterogeneity revealed by sampling and natural glioma evolution over time. We did not detect any evidence for mouse specific evolution. Minor changes in clonal trajectories have also been observed in certain PDX from GBMs^67^, brain metastases^86^ and other cancers^31,87^. In this respect, PDOX models can be considered as a proxy for dynamic clonal evolution, which is difficult to measure in patients. We also did not observe major changes in paired longitudinal glioma samples neither in the parental patient tumor nor in the corresponding PDOX, in accordance with limited treatment-induced clonal evolution in diffuse gliomas^18^. We report a case of clonal evolution from *EGFRvII to EGFRvIII*, which was recapitulated in the corresponding PDOXs. Although *EGFRvIII* may be lost upon recurrence, cases with acquisition of this variant were also reported^17,88^. Interestingly, longitudinal models also retained state-specific intra-tumoral heterogeneity and genetic subclones, highlighting the notion that these unique matched PDOXs provide an ideal platform to study specific molecular events in initial and recurrent disease side by side. We further show that propagation of GBM cells grown as GSCs *in vitro* leads to faster genetic drift, including ploidy changes and acquisition of new CNAs and genetic variants.

At scale tumor-intrinsic epigenetic and transcriptomic profiles of individual tumors were well recapitulated in PDOX. Our PDOX cohort represents diverse molecular subtypes and retains intra-tumoral heterogeneity and plasticity, in particular, we find that GBM PDOX display cellular state transitions recently described in patient samples^58^. No major molecular changes or selection of cellular subpopulations were detected, except for those related to the replacement of human TME by mouse counterparts. These changes are expected in bulk tissue analyses where methylation and transcriptome profiles are biased by TME-derived signals^5,53^. In line with previous reports^26,27^ *in vitro* cell lines showed increased global DNA methylation levels and more profound changes in transcriptomic profiles.

Limitations of PDOX models include possible interspecies differences at the molecular level and the lack of a complete immune system in immunocompromised animals. While the adaptive immune system is largely absent in these mice, they retain a largely functional innate immune system, including microglia, the brain resident immune effector cells, and peripheral myeloid cells. GBM are largely lymphocyte depleted tumors^89^, while microglia and macrophages constitute the major immune component^90^. Here we show that classical glioma TME components such as microglia/macrophages, astrocytes and OPCs are present in xenografted tumors, indicating that tumor cell interactions with the TME remain active in PDOX. Of note, we observe a similar transcriptomic shift in tumor-associated microglia/macrophages as described in GBM patients^59,91^. It remains to be determined to what extent these models will be amenable to immunotherapeutic studies targeting tumor-associated microglia/macrophages. Though challenging, adaptation of glioma PDOXs to a humanized background might be possible and/or studies in an immunocompetent context could be performed ex vivo with PDOX-derived 3D organoids co-cultured with autologous immune cells.

Other drawbacks of patient-derived (orthotopic) xenografts, include high costs, complex logistics and an inherent low-throughput nature. Large-scale *in vivo* screens are possible; however, they are laborious and require specific statistical settings^31^. Expansion of human gliomas in PDOX and initial drug screens performed on PDOX-derived organoids appears as a good compromise between retention of glioma hallmarks and a cost-effective drug testing pipeline. In contrast to patient-derived short-term cultures and organoids^29,63^, it allows for tumor expansion and *in vivo* validation. We have developed our protocols to reconstitute 3D organoids of equal size, which allowed for reproducible drug testing. Downscaling of cell number per organoid facilitated drug delivery, viability detection and upscaling to high-throughput screens. These protocols can also be adapted to reintroduce TME components^44^ and immune cells. We show that PDOX-derived organoids show clinically relevant responses: (i) organoids with *MGMT* promoter methylation showed higher sensitivity to TMZ, (ii) *CDK4/6* amplified organoids responded better to CDK4/6 inhibitors, (iii) organoids carrying *EGFR R108K, G598V* and *A289T* gain-of-function mutations were most sensitive to Erlotinib, whereas EGFR low tumors were most resistant. Although EGFRvIII^92^ and deletions in the C-terminal domain (Δ25-27/28) were shown to sensitize GBM cells to Erlotinib^93^, none of the EGFR structural variants present in our testing group systematically sensitized tumors to any of the EGFR inhibitors. Of note, the tested organoids displayed *PTEN* loss, a known resistance factor leading to dissociation of EGFR inhibition from downstream PI3K pathway inhibition^92^. Remarkably, Dianhydrogalactitol (DAG, VAL-083) showed a significantly better response than TMZ against GBMs of different genetic backgrounds and irrespective of *MGMT* status. DAG’s efficacy was confirmed *in vivo*, with no toxicity observed, lending optimism to ongoing clinical trials.

Overall, our glioma PDOX cohort provides a useful platform for understanding tumor biology and preclinical treatment interventions at the individual patient level. Although the parallel use of PDOXs as patient avatars during treatment remains challenging due to the poor prognosis of GBM patients, PDOXs can play a key role in ‘mouse clinical trials’^94^ for personalized medicine regimens. Longitudinal models will further serve as a powerful tool for analysis of tumor evolution and resistance mechanisms upon general and targeted therapies. By sharing the models and molecular data we aim to facilitate large collaborative preclinical trials in the future.

## MATERIALS AND METHODS

### Clinical samples and PDOX derivation

Glioma samples were collected at the Centre Hospitalier of Luxembourg (CHL; Neurosurgical Department) from patients having given informed consent, and with approval from the local research ethics committee (National Committee for Ethics in Research (CNER) Luxembourg). Samples P8, P13 and P3 were obtained from Haukeland University Hospital (Bergen, Norway) following approval of the local ethics committee. Small pieces of tissue were flash frozen for further molecular analysis. If enough tumor material was obtained, 3D organoids from patient samples were prepared as previously described^38^. Briefly, mechanically minced fresh human glioma tissue pieces, without enzymatic digestion, were seeded on agar coated flasks (0.85%) and allowed to form organotypic spheroids (here called organoids) for up to 2 weeks at 37°C under 5% CO2 and atmospheric oxygen in DMEM medium, 10% FBS, 2 mM L-Glutamine, 0.4 mM NEAA and 100 U/ml Pen-Strep (all from Lonza). Organoids (generation 0) with a diameter of 300-1000 μm were then implanted in the brain of immunodeficient mice (NOD/Scid or NSG; 6 organoids per mice) using a Hamilton syringe (Hamilton, Reno, NV, USA). Where indicated organoids were implanted into the brain of nude rats (rnu-lrnu-; 10 organoids per rat: P3, P8, P13 models). Animals (generation 1) were maintained under SPF conditions and sacrificed at the appearance of neurological (locomotor problems, uncontrolled movements) or behavioral abnormalities (prostration, hyperactivity) and weight loss. Optionally tumor volume was monitored by MRI. Organoids (generation 1) were further prepared from minced xenografted brains as for patient tissue and serially implanted for several generations. A PDOX model was considered to be established at generation 3, when tumor phenotype and animal survival appeared stable. For specific purposes, experiments were performed in nude mice and/or eGFP expressing NOD/Scid mice^95^. Kaplan-Meier survival curves were produced in GraphPad with Wilcoxon signed-rank statistical test. The handling of animals and the surgical procedures were performed in accordance with the regulations of the European Directive on animal experimentation (2010/63/EU) and the Norwegian Animal Act, *i.e*., the experimental protocols were approved by the local ethics committee (Animal Welfare Structure of the Luxembourg Institute of Health; protocols LRNO-2014-01, LUPA2019/93 and LRNO-2016-01) and by the Luxembourg Ministries of Agriculture and of Health. PDOX models are available from the corresponding author or via EuroPDX consortium (https://www.europdx.eu/) and PDXFinder (https://www.pdxfinder.org/).

### Magnetic Resonance Imaging

During image acquisition mice were kept under anesthesia with 2.5% of isoflurane, with constant monitoring of breathing and temperature. For routine follow up, mice were placed in the MRI (3T MR Solutions) and a Fast Spin Echo T2-weighted MRI sequence was applied, with field of view of 25 mm, matrix size of 256×256, TE of 68ms, TR of 3000ms and slice thickness of 1 mm. To visualize the contrast enhancement, T1-weighted sequences without and with contrast injection were used. Fast Spin Echo T1-weighted MRI was defined with the following parameters: field of view of 25 mm, matrix size of 256×252, TE of 11ms, TR of 1000ms and slide thickness of 1 mm. Contrast agent (Gadodiamide, Omniscan, GE-Healthcare) at 0.5mmol/kg was injected intravenously 1min prior to the scan. MRI data was analyzed by ImageJ.

### Cell lines and cell line-derived xenografts

Glioma stem-like cell (GSC) cultures (P3NS, P13NS, T16NS, T158NS, T226NS, T384NS, T394NS, T407NS) were derived from PDOXs by papain-based enzymatic digestion of PDOX tissue and cultured as 3D spheres in serum-free medium based on Neurobasal^®^ base medium (Life Technologies) supplemented with 1 x B27 (Life Technologies) 2 mM L-Glutamine, 30 U/ml Pen-Strep, 1 U/ml Heparin (Sigma), 20 ng/ml bFGF (Miltenyi, 130-093-841) and 20 ng/ml EGF (Provitro, 1325950500). GSC NCH601, NCH421k and NCH644 lines^24^ were cultured as non-adherent spheres in DMEM-F12 medium (Lonza) containing 1 x BIT100 (Provitro), 2 mM L-Glutamine, 30 U/ml Pen-Strep, 1 U/ml Heparin (Sigma), 20 ng/ml bFGF (Miltenyi, 130-093-841) and 20 ng/ml EGF (Provitro, 1325950500). U87 and U251 cells (obtained from ATCC, HTB-14) were cultured as adherent monolayers in DMEM containing 10% FBS, 2 mM L-Glutamine and 100 U/ml Pen-Strep (all from Lonza). Cell lines were regularly tested for mycoplasma contamination. Cell lines were authenticated by DNA profiling using a SNP-based multiplex approach and compared to the other continuous cell lines in the DSMZ database. SNP profiles were unique. For *in vivo* experiments tumor cells (50’000-100’000 per mouse) were slowly injected through a Hamilton syringe (Hamilton, Reno, NV, USA) into the right frontal cortex. The animals were sacrificed upon weight loss, appearance of severe neurological (locomotor problems, uncontrolled movements) or behavioral abnormalities (prostration, hyperactivity).

### Immunohistochemistry and neuropathological analysis

Coronal sections from paraffin-embedded brains were stained with hematoxylin (Dako) and 1% eosin (H&E) (Sigma). For immunostaining, sections were pre-treated for 5min with Proteinase K (Dako) followed by 30 min incubation at 95°C in retrieval solution (Dako). The Dako Envision+System-HRP was used following the manufacturer’s instructions. Primary and secondary antibodies were incubated for 1h. Signal was developed with 3,3’-diaminobenzidine chromogen in 5–20 min. Additional IHC preparations were performed using a Discovery XT automated staining module (Ventana) and standard protocols (list of antibodies in **Table S7)**. The existence of necrosis and the degree of invasion was assessed on the basis of H&E and human-specific Nestin staining. Proliferation index was determined as % Ki67-positive cells per whole cell population. An index of 37% was used to split Ki67 low and high models. IHC of mouse endothelial cells (CD31) was performed on isopentane flash-frozen tissues and cryostat sections (10μm) were fixed with acetone and chloroform. Nonspecific binding was blocked with 2% FBS in TBS and antibodies were incubated for 1h at RT. Pictures were acquired with a Leica DMI 6000B microscope. Vessel quantification was done using ImageJ software. Average vessel area (μm^2^) was used as a proxy for vessel abnormality. Vessel area high and low models were analyzed after median split dichotomization into two groups. Kaplan-Meier survival analyses, including Log-rank and Wilcoxon testing were performed in GraphPad Prism 8. Other analyses were performed with two-tailed Student’s t-test.

### Flow cytometry

Tumor and PDOX tissue was dissociated with MACS Neural Tissue Dissociation Kit (P) (Miltenyi) following manufacturers’ instructions. For phenotyping flow experiments were performed as described^55^. Single cell suspensions were resuspended in DMEM, containing 2% FBS, 10 mM HEPES pH 7.4 and DNase I (10 μg/ml; Sigma) at 1×10^6^ cells/ml followed by 90 min incubation with Hoechst 33342 (5 μg/ml, Bisbenzimide, Ho342; Sigma) at 37°C. After washing, cells were resuspended in ice-cold HBSS 2% FBS, 10 mM HEPES pH 7.4 buffer (100 μl/test). Prior to flow cytometric analysis, cells were incubated with the IR-LIVE/DEAD^®^ Fixable Dead Cell Stains (Invitrogen; 1 μg/ml) and appropriate preconjugated antibodies for 30 min at 4°C in the dark (**Table S7**). Data acquisition was performed on a FACS Aria^™^ SORP cytometer (BD Biosciences) fitted with a 632 nm (30 mW) red laser, a 355 (60 mW) UV laser, a 405 nm (50 mW) violet laser and a 488 nm (100 mW) blue laser was used. Data analyses were done with DIVA software (BD Bioscience). For cell sorting, single cell suspensions were stained with the TO-PRO^®^-3 shortly before sort. eGFP-negative tumor cells and eGFP-positive mouse non-neoplastic cells were sorted to cold flow cytometry buffer, centrifuged and resuspended in organoid culture medium. Organoids free of non-neoplastic cells were obtained from sorted eGFP-negative tumor cells by plating 20000 cells per well of 24-well plates pre-coated with agar. For mixed organoids 20000 sorted tumor cells were premixed with 2000 sorted eGFP-positive non-neoplastic mouse cells (10%). Alternative, FACS-sorted GFP-negative tumor cells were washed in cold HBSS buffer and processed directly to RNA extraction.

### Ploidy assessment

Nuclei were isolated from liquid nitrogen flash frozen PDOX tumors^39^. Samples were minced in DAPI buffer [10 μg/ml DAPI in 146 mM NaCl, 10 mM Tris-HCl (pH 7.5), 0.2% IPEGAL]. Nuclei were disaggregated subsequently with 20G and 25G needles and filtered through a 50 μm and a 30 μm mesh. Tumor nuclei were stained with the human-specific anti-Lamin A/C-PE antibody (**Table S7**). Optionally, PDOX-derived single cell tumor cells and cell lines were stained with IR-LIVE/DEAD^®^ Fixable Dead Cell Stains (Invitrogen; 1 μg/ml) and fixed with cold 80% ethanol. PBMCs were added to each sample as internal diploid control. Flow analysis was carried out with Aria^™^SORP or Canto^™^ flow cytometers (BD Biosystems). DNA content was analyzed with the FlowJo software.

### Extraction and quality control of genomic DNA

DNA from flash frozen primary patient tissue, PDOX tumor tissue, PDOX-derived organoids and GSC cultures was extracted using the AllPrep DNA/RNA Mini Kit^®^ (Qiagen) following manufacturer’s instructions for “Simultaneous purification of genomic DNA and total RNA from animal tissues”. DNA was eluted in 50 μl of Nuclease-free water and concentrations were measured using a NanoDrop 1000 (Thermo Fisher Scientific). Integrity of gDNA was analyzed with a 1 % E-Gel^™^ EX Agarose Gel (Thermo Fisher Scientific). To obtain DNA from formalin-fixed, paraffin-embedded (FFPE) samples, the tissue block was punched to obtain a tissue core of 2 mm containing at least 70% tumor tissue. After a deparaffinization step (Deparaffinization solution, Qiagen), DNA extraction was performed using QiAamp DNA FFPE tissue kit (Qiagen) according to the manufacturer’s instruction. DNA concentrations were measured on the Qubit 4.0 fluorometer (Thermo Fisher Scientific), using the Qubit dsDNA BR Assay kit (Thermo Fisher Scientific).

### Array comparative genomic hybridization (array-CGH)

Array-CGHs were performed as previously described^39^ with the following changes. DNA was fragmented (200-500bp) using enzymatic digestion with RSA1 and Alu1 (Agilent Technologies) and labeled with the BioPrime array-CGH Genomic labeling Kit (Life Technologies) and Cy3 and Cy5 dyes (GE Healthcare) following standard protocols for Agilent array-CGH (CGH enzymatic protocol v6.2; Ref # G4410-90010). Female or Male gDNA pool (Promega) was used as a reference. All labelling reactions were assessed using a Nanodrop 1000 (Thermo Fisher Scientific) before mixing and hybridized to either a 1×1M, 2×400K, 4×180K or 8×60K SurePrint G3 human CGH microarray (Agilent Technologies) according to manufacturer’s instructions (CGH enzymatic protocol v6.2; Ref # G4410-90010). Microarray slides were scanned using an Agilent 2565C DNA scanner and images were analyzed with Agilent Feature Extraction version 12.5, using default settings. Data was assessed with a series of quality control metrics and analyzed using an aberration detection (ADM2) implemented in the CytoGenomics software versions 4.2 and 5.0.2.5 (Agilent Technologies). Aberrations were called using the ADM2 algorithm with a threshold setting of 6 and an aberration filter with a minimal number of probes = 3 and a minimal AvgAbsLogRatio = 0.25. For correlation analysis, each sample was initially processed with *Agilent CytoGenomics 4.2* in order to obtain the characterization of genomics regions (BED files) described as one of the following events: “amplification”, “gain”, “loss” or “deletion. Next, from each file, only regions > 50kb were extracted in order to construct a reference mapping file using a combination of ‘*intersectBed’* and *‘multiIntersectBed’* functions from the BEDtools suite. Finally, BED files were mapped on that common reference with their corresponding type of event. As a consequence the resulting matrix represents features detectable by any of the four array types. Chromosomes X and Y were removed. Hierarchical clustering showed no bias arising from the array type used. Pearson correlation was applied to assess relationships between genetic profiles of each sample. Next, we estimated the effects of the experimental factors on DNA copy number variation data. As these data were represented by integer values between −2 and 2, we were unable to fit a global linear model. Instead, we used a chi-squared contingency table test implemented in the ‘stat’ package of R. Independently for each factor and for each DNA site we tested, whether a distribution of copy numbers is different for different factor levels of the corresponding factor. Mean −log10(p-value) and mean chi-squared statistics were reported for graphical presentation.

### Targeted DNA sequencing

500 ng of extracted gDNA were diluted in 130 μl low TE buffer (Qiagen) and sheared via sonication on a Bioruptor^®^ UCD-200 (Diagenode) to an average fragment size of 150-300 bp. DNA fragment size was determined using the DNA 1000 Kit on the Bioanalyzer 2100 (Agilent Technologies). A custom-made Agilent SureSelect^XT^ Target Enrichment Library (Cat No. G9612B) was used for Illumina Paired-End Multiplexed Sequencing on a MiSeq^®^ instrument (Illumina). The panel design 1 for the Target Enrichment Library was fully adapted from^47^ (181 genes and 3 promoters). Further design changes were made using SureDesign - Agilent eArray (Agilent Technologies) to produce the panel design 2, containing additional regions (234 genes and 3 promoters). A total of 59 samples were sequenced 22 samples with the panel 1 and 37 samples with the panel 2. Library preparation was performed according to manufacturers’ instruction. The Illumina MiSeq^®^ Reagent Kit v3 (Cat No. MS-102-3003) was selected applying the Illumina reagent selection algorithm (https://emea.support.illumina.com/downloads/sequencing_coverage_calculator.html).

Variant calling was done as follows: Raw sequencing reads (fastq) were quality trimmed using the tool fastp (v. 0.20.0)^96^. Trimmed reads were aligned to an *in silico* fused reference genome (ICRG) containing the human genome GRCh37.75 (ENSEMBL) and the mouse genome mm10 using BWA mem (v. 0.7.17)^97^. Reads that mapped to human chromosomes were extracted from the bam file using SAMtools (v.1.9) and realigned to the human reference genome only^98^. Duplicates were annotated and removed using MarkDuplicates under GATK (v.4.0.5.1). Bam statistics were assessed using SAMtools and compared between the initial mapping to the ICRG, the realignment to the human genome and after removing duplicates. Single nucleotide variants (SNVs) and smaller insertions and/or deletions (indels) were called in the CLC Genomics Workbench (v.12.0.3) using deduplicated mappings. Variants were only called in regions with a minimum coverage of 10 reads and a minimal allele frequency of 5 %. All variants that were likely to be polymorphisms and occurred in more than 1% of the gnomAD (v.2.0.2) data base were filtered out. SNVs were annotated with COSMIC (v.89), ClinVar and dbSNP (v.150)^99^. The primary focus in SNV calling was to determine coding changes (missense and inframe mutations), truncating (stop and frameshift mutations) and splice site mutations. Due to poor coverage of *TERT* promoter, this region was excluded from the global analysis. All filtered variants were manually checked to exclude artefacts and variants were further classified according to the American College of Medical Genetics and Genomics (ACMG)^100^. Only pathogenic, likely pathogenic or variants of uncertain significance (VUS) were reported. For comparing the %-overlap of variants between patient tissues and PDOX, the following thresholds were applied: Minimum coverage 10, Minimum count 2, Minimum frequency 5 %. Reads were mapped with a linear and an affine gap cost mapping and variants were merged after calling from both mappings.

Structural variants (SVs) and copy number alterations (CNAs) were analyzed using Manta (v. 1.6.0) and CNVkit (v.0.9.6)^101,102^. For these analyzes alignment files with marked duplicates were used. SVs were annotated with SnpEff (v. 4.3.1t)^103^ and filtered for variants with at least 5 supporting paired and/or split reads. CNAs were called in two separate groups, as two versions of the sequencing panels were used and the target region is important for CNA calling via CNVkit. For panel 1 no reference samples were sequenced and CNA calling was performed against a flat reference. CNAs of all samples that were sequenced with the panel 2 were normalized against a reference created from normal samples including the commercial available male (Cat No. G1471) or female (Cat No. G1521) references from Promega (Madison, Wisconsin, US) and DNA from blood of two patients. Segmentation was performed using circular binary segmentation according to default settings. Gene metrics were determined for all variants with a minimum log2 deviation of 0.4.

Workflow automation was performed using the workflow manager snakemake (v.5.6.0) under conda (v.4.7.12)^104^. Additional data handling was performed applying R (v.3.6) in the environment of RStudio (v.1.1.456). All CNAs and the SVs in EGFR were visualised, manually checked and compared to available data from array-CGH and array-based DNA methylation analysis.

Subclonal deconvolution via PyClone was performed in parallel with the above data in an independent manner. PyClone input requires variants and copy number. To acquire these data, reads were aligned to hg38, processed with Picard’s MarkDuplicates {http://broadinstitute.github.io/picard/}, and GATK indel realignment and base recalibration performed^105^. Variants were called using mpileup (Samtools v.1.9)^98^ and Varscan 2’s (v.2.4.4) pileup2snp and mpileup2indel commands^98^ with default settings but a p-value of 1.00. Only positions in targeted regions were kept. Variants in dbSNP were filtered out. Absolute copy numbers were estimated using array-CGH. Log2 ratios were segmented using DNAcopy (v1.52.0)^106^. A custom script estimated purities and absolute copy numbers based on the assumption that chr7 likely had a clonal single copy gain, resulting in inference of one copy loss of chr10 and one copy gains of chr19 and chr20 (common events in GBM) in all analysed samples (T192, T233, T251, T158, T347, T470), validating this approach. PyClone (v.0.13.1)^107^ was run under default settings, with the addition of ‘--prior total_copy_number’ to indicate the use of total copy numbers. Purities were taken from the array-CGH estimates for biopsies, and was set to 1.00 for PDOX samples.

### Digital PCR

Digital PCR was used to detect and quantify IDH1 R132H in genomic DNA using QuantStudio 3D Digital PCR System and IDH1 Digital PCR Mutation Detection Assays from Thermo Fisher (Assay ID # Hs000000036_rm for c.395G>A (p. R132H)) according to the manufacturer’s instructions. The reaction volume was 14.5 μL containing 7.5 μl QuantStudio 3D Digital PCR Master Mix v2 (Thermo Fisher Scientific, cat#: A26359), 0.73 μl of assay and sample DNA. Each assay contained forward and reverse primers, and 2 specific dye-labeled probes. The first one with a Vic reporter dye linked to the 5’end and an MGB linked to the 3’ end to detect the WT allele. The second one with a FAM reporter dye at the 5’end and an MGB at the 3’ end to detect the mutant allele. The thermal cycling conditions were 96 °C for 10 min; 39 cycles of 60 °C for 2 min and 98 °C for 30 sec; final extension at 60 °C for 2 min. Two replicates of each sample were run and DNA input amount was 20 ng per chip. Human Genomic DNA Male (Promega, cat # G1471) and IDH1 R132H Reference Standard (Horizon, cat # HD677) were used as wild type reference DNA and positive reference respectively. Data analysis was done with the QuantStudio 3D Analysis Suite Cloud Software version 3.1.5; chips with <15000 partitions above the default quality threshold were omitted.

### Array-based DNA methylation Analysis with Infinium^®^ MethylationEPIC

Processing of HumanMethylationEPIC BeadChip arrays was performed by the Helmholtz Zentrum Muenchen (Research Unit of Molecular Epidemiology/Institute of Epidemiology, German Research Center for Environmental Health, Neuherberg, Germany)^108^ or at the Laboratoire National de Santé in Luxembourg. Bisulfite conversion of 250-500 ng of gDNA was done using the EZ DNA Methylation Kit (Zymo Research) according to manufacturer’s procedure, with the alternative incubation conditions recommended when using the Illumina Infinium^®^ Methylation Assay. After bisulfite treatment, Infinium HD FFPE Restore kit (Illumina) protocol was performed on 8 μl of DNA from FFPE samples. Genomewide DNA methylation was assessed using the HumanMethylationEPIC BeadChip (Cat No. WG-317-1001), following the Illumina Infinium^®^ HD Methylation protocol. This consisted of a whole genome amplification step using 4 μl and 8 μl (for fresh-frozen and FFPE samples, respectively) of each bisulfite converted sample, followed by enzymatic fragmentation and hybridization of the samples to BeadChips (Illumina). After a step of single-nucleotide extension, the BeadChips were fluorescently stained and scanned with Illumina HiScan SQ scanner or iScan System. Additional Illumina HumanMethylation450 BeadChips were processed according to manufacturer’s instruction at the German Cancer Research Center (DFKZ) Genomics and Proteomics Core Facility. Raw scanning data was normalized using GenomeStudio (Illumina) to obtain beta-values (CpG methylation estimates). Raw Intensity Data files (.idat) were exported from the BeadArray. Pearson correlation was applied to assess relationships between epigenetic profiles of each sample. The R package *‘RnBeads’* was used to generate individual 450k and EPIC RnBeadSets^109^ that were normalized using the *‘BMIQ’* method^110^. Both platforms were combined using the *‘rnb.combine.arrays’* function in order to extract only common sites present in both objects with corresponding DNA methylation level. The DNA methylation level for each locus was measured as a beta-value score; that can range from zero to one with scores of zero indicating completely unmethylated DNA and scores of one indicating complete methylated DNA. Hierarchical clustering showed no bias arising from the array type used. Pearson correlation was applied to assess relationships between genetic profiles of each sample.

As several of the considered factors were strongly correlated, we estimated their importance by consequent fitting unavailable ANOVA models, independently for each CpG site and factor. Mean F-statistics over all variable CpG cites was then used to illustrate the importance of the factors. To detect differentially methylated regions (DMRs) or CpGs (DMCs), IDAT files were subjected to background correction, global dye-bias normalization, calculation of DNA methylation level, and detection p-values using *‘methylumi.noob’* within the *‘RnBeads’* package. Differential methylation analysis was conducted on genomic site and region level according to sample groups (Patient vs. PDOX or IDHwt vs. IDH1mut) using *‘limma’* and fitted using an empirical Bayes approach on M-values^111^. In general, array probes were divided into 4 different genomic regions, giving info on functional genomic distribution: 1) Tiling regions with a window size of 5kb distributed over the whole genome, 2) Genes and 3) Promoters annotated with Ensembl gene definitions from the biomaRt package. Promoters were defined as the region spanning 1,500 bases upstream and 500 bases downstream of the TSS of the corresponding gene. 4) CpG islands tracked from UCSC genome browser. Furthermore, probes were divided into those within CpG islands (CGI), in CGI shores, shelves, or open seas (with or without overlapping gene bodies). In the comparison between ‘Patient’ and ‘PDOX’, the following criteria were selected: adj. p-value < 0.01, absolute difference in mean methylation β value > 0.2. Beta value distribution plots for probe categories (‘Open Sea’, ‘Shelf’, ‘Shore’ or ‘Island’) were extracted from the integrated ‘Exploratory Module’ from *‘RnBeads’*. Global beta value density plots for longitudinal samples were generated using the *‘minfi’* package in R, after Noob background correction and global dye-bias normalization. Analysis of CpG methylation signatures was performed as described previously^14^. DNA methylation-based classification was performed at https://www.molecularneuropathology.org/mnp as described previously^48^.

### Genome-wide gene expression analysis

Total RNA was extracted using the QIAGEN^®^ RNeasy Mini Kit according to the manufacturer’s protocol. GeneChip^®^ Human Gene 1.0ST Arrays were used to determine the expression profiles. Total RNAs were processed using the Ambion WT expression kit (Life Techniologies) and the Affymetrix WT Terminal & Labeling kit before being hybridized on Affymetrix arrays according to the manufacturer’s instructions (protocol P/N 702808 Rev.6). Upon hybridization, microarrays were washed, stained and scanned according to manufacturer’s standard procedures. Affymetrix CEL files containing hybridization raw signal intensities were processed to gene expression signals using the RMA (robust multichip average) algorithm implemented in the *oligo* package (version 1.44.0). *hugene10sttranscriptcluster.db* package version 8.7.0 was then used to map Affymetrix ID to entrez gene ID. R statistical environment was used for hierarchical clustering, principal component analysis and for empirical Bayesian statistics (LIMMA^111^, R/Bioconductor). List of differentially expressed genes (DEG) were obtained with the eBayes/LIMMA. FDR was calculated with the Benjamini and Hochberg approach^112^, Thresholds were set up for FDR<0.01 and absolute fold change (abs(FC))≥2. The Metascape^®^ database^113^ was used for data mining.

The similarity between our patient biopsies, PDOXs and cell lines with GBM tumors from The Cancer Genome Atlas (TCGA) cohort (538 GBM samples) was investigated using gene expression data^54^. Our cohort’s data were ranked based on their interquartile ranges to select the top-5000 (most variable) probes across samples. We focused on probes with mapped gene symbols, for genes with multiple probes their expression values were (mean) merged with Babelomics 5^114^. Filtering resulted in 4069 unique gene symbols. TCGA data were downloaded from The Broad Institute GDAC Firehose (http://gdac.broadinstitute.org), and the pre-processed gene expression data (RSEM values) were analyzed. Gene symbols from our cohort were matched to the TCGA data, and 2420 unique symbols were unambiguously found in both datasets. Using the expression data for these genes, we measured (Spearman) correlation coefficients between our cohort samples and TCGA tumors. The resulting correlations with the TCGA tumors were ranked and graphically visualized in terms of individual samples and sample groups. Analyses were implemented with the R statistical language, packages corrplot and ggplot2 (https://www.r-project.org).

Consensus independent component analysis (ICA), a data-driven dimensionality reduction method that performs a matrix decomposition, was applied to assess signals arising from non-malignant cells. ICA with *k* components represents log2-transformed gene expression matrix ***X*** as a matrix product of matrices ***S*** (signals) and ***M*** (weights). The first shows contribution of genes in *k* statistically independent signals. Biological meaning of these signals was detected by functional annotation of the most contributing genes. In order to improve reproducibility of ICA decomposition, which can be affected by the selection of initial estimations, we applied consensus ICA approach^115^. ICA was run multiple times and the resulted matrices ***S*** and ***M*** were mapped and averaged between the runs. The analysis of the cell lines and TCGA reference dataset was performed as described in^115^.

### Single cell RNA-Seq using Drop-Seq

For scRNA-seq experiments PDOXs derived in nude mice were used. To obtain a pure population of single viable cells and to distinguish human tumor cells from mouse TME subpopulations PDOXs were dissociated and FACS-sorted (P3, P8, P13) or MACS-purified (T16, P13). For FACS we have separated hCD90 positive tumor cells from hCD90 negative mouse TME subpopulations^56^. MACS-based purification was performed with *Myelin Removal* Beads II followed by Mouse Cell Depletion kit (Miltenyi Biotec) according to manufacturer’s protocols. Prior to cell loading on the Drop-seq chips, the viability of cells was verified and concentration was adjusted to ~150 cells/μl as optimal concentration to achieve single cell encapsulation within each droplet of ~1 nl. All samples analyzed had a cell viability > 95%.

Microfluidics devices were fabricated using a previously published design^116^. Soft lithography was performed using SU-8 2050 photoresist (MicroChem) on 4” silicon substrate to obtain a feature aspect depth of 100 μm. After overnight silanization (using Chlorotrimethylsilane, Sigma), the wafer masks were used for microfluidics fabrication. Drop-seq chips were fabricated using silicon based polymerization chemistry. Briefly, Polydimethylsiloxane (PDMS) base and crosslinker (Dow Corning), were mixed at a 10:1 ratio, mixed and degassed before pouring onto the Drop-seq master template. PDMS was cured on the master template, at 80°C for 2h. After incubation and cooling, PDMS slabs were cut and the inlet/outlet ports were punched with 1.25 mm biopsy punchers (World Precision Instruments). The PDMS monolith was plasma-bonded to a clean microscopic glass slide using a Harrick plasma cleaner. Immediately after pairing the plasma-treated surfaces of the PDMS monolith and the glass slide, flow channels of the Drop-seq chip were subjected to a hydrophobicity treatment using 1H,1H,2H,2H-Perfluorodecyltrichlorosilane (in 2% v/v in FC-40 oil; Alfa Aeser/Sigma). After 5 min of treatment, excessive silane was blown through the inlet/outlet ports. Chips were further incubated at 80°C for 15 min.

Experiments followed the original Drop-seq protocol^116^ with minor changes. Synthesized barcoded beads (Chemgenes corp., USA) were co-encapsulated with cells inside the droplets containing lysis reagents using an optimal bead concentration of 200 beads μl^-1 in Drop-seq Lysis buffer medium. Cellular mRNA was captured on the beads via barcoded oligo (dT) handles synthesized on the surface. For cell encapsulation, 2 ml of cell and bead suspensions were loaded into 3 ml syringes (BD), respectively. To keep beads in homogenous suspension a micro-stirrer was used (VP scientific). The QX 200 carrier oil (Bio-rad) used as continuous phase in the droplet generation was loaded into a 20 ml syringe (BD). For droplet generation, 3.6 ml per h and 13 ml per h were used in KD scientific Legato syringe pumps for the dispersed and continuous phase flows, respectively. After stabilization of droplet formation, the droplet suspension was collected into a 50 ml Falcon tube. Collection of the emulsion was carried out until 1 μl of the single cell suspension was dispensed. Droplet consistency and stability were evaluated by bright-field microscopy using INCYTO C-Chip Disposable Hemacytometer (Fisher Scientific). Bead occupancy within droplets was carefully monitored to avoid multiple bead occupancy. The subsequent steps of droplet breakage, bead harvesting, reverse transcription and exonuclease treatment were carried out in accordance to^116^. RT buffer contained 1x Maxima RT buffer, 4 % Ficoll

PM-400 (Sigma), 1 μM dNTPs (ThermoScientific), 1 U/ml Rnase Inhibitor (Lucigen), 2.5 μM Template Switch Oligo, and 10 U/ml Maxima H-RT (ThermoScientific). Post Exo-I treatment, the bead counts were estimated using INCYTO C-Chip Disposable Hemacytometer, and 10,000 beads were aliquoted in 0.2 ml Eppendorf PCR tubes. PCR mix was dispensed in a volume of 50 μl using 1x Hifi HotStart Readymix (Kapa Biosystems) and 0.8 mM Template-Switch-PCR primer. The thermocycling program for the PCR amplification was modified for the final PCR cycles by 95°C (3 min), four cycles of 98°C (20s), 65°C (45s), 72°C (3 min), 10 cycles of 98°C (20s), 67°C (20s), 72°C (3 min) and followed by a final extension step of 72°C for 5 min. Post PCR amplification, libraries were purified with 0.6x Agencourt AMPure XP beads (Beckman Coulter), in accordance with the manufacturer’s protocol. Finally, the purified libraries were eluted in 20 μl RNAse/DNAse-free molecular grade water. Quality and concentration of the sequencing libraries were assessed using BioAnalyzer High Sensitivity Chip (Agilent Technologies). The 3’ end enriched cDNA libraries were prepared by tagmentation reaction of 600 pg cDNA library using the standard Nextera XT tagmentation kit (Illumina). Reactions were performed according to the manufacturer’s instructions, samples were barcoded using the N7xx index series and 400 nM custom P5 hybrid primer: (AATGATACGGCGACCACCGAGATCTACACGCCTGTCCGCGGAAGCAGTGGTATCAACGCAGAG T*A*C). The PCR amplification cycling program used was: 95°C 30s; fourteen cycles of: 95°C (10s), 55°C (30s), 720°C (30s) followed by a final extension step of 72°C (5 min). Libraries were purified twice to reduce primers and short DNA fragments with 0.6x and 1x Agencourt AMPure XP beads (Beckman Coulter), respectively, in accordance with the manufacturer’s protocol. Finally, purified libraries were eluted in 15 μl molecular grade water. Quality and quantity of the tagmented cDNA library was evaluated using BioAnalyzer High Sensitivity DNA Chip. The average size of the tagmented libraries prior to sequencing was between 400-700 bps.

Purified Drop-seq cDNA libraries were sequenced using Illumina NextSeq 500 with the recommended sequencing protocol except for 6pM of custom primer (GCCTGTCCGCGGAAGCAGTGGTATCAACGCAGAGTAC) applied for priming of read 1. Paired end sequencing was performed with the read 1 of 20 bases (covering the random cell barcode 1-12 bases and the rest 13-20 bases of random unique molecular identifier (UMI) and for read 2 the 50 bases of the genes.

Bioinformatic processing followed the DropSeq protocol^116^ using the DropSeq tool version 1.16. In brief, FASTQ files were assembled from the raw BCL files using Illumina’s bcl2fastq converter and ran through the FASTQC codes [Babraham bioinformatics; https://www.bioinformatics.babraham.ac.uk/projects/fastqc/] to check library qualities by the assessment parameters a) quality per base sequence, b) per base N content, c) per base sequence content and d) over-represented sequences. Libraries with significant deviation were re-sequenced. FASTQ files were subsequently merged and converted to binaries using PICARD’s fastqtosam algorithm. The resulting digital gene expression matrix (DGE) was first cut based on knee plot analysis and subsequently filtered by the Seurat version 3 and Monocle version 2 packages (http://cole-trapnell-lab.github.io/monocle-release/) in R (version 3.6.0) based on ribosomal and mitochondrial genes as well as on low transcript content. The following threshold filters were used: only cells that expressed at least 200 genes and presented 1×10^6^ total mRNAs, and only genes which were expressed in at least 5 cells were considered for further analysis. To normalize for transcript capturing between the beads, the averaged normalized expression levels (log2(TPM+1)) were calculated. After filtering and normalization, our dataset included 3138 cells (per sample cell counts: P3 =543 cells, P8= 502 cells, P13= 1295, T16=798 cells). To examine relative expression levels, we centered the data by subtracting the average expression of each gene from all cells. Digital gene expression matrix of the TME subpopulations of PDOX P8 and normal mouse brain was filtered and normalised as described above. After filtering and normalisation, the dataset included 892 cells (per sample cell counts: P8 = 453 cells, Control= 439 cells). Dimensionality reduction and gene expression markers identification and visualisation were done using UMAP implemented in the Seurat package version 3^117,118^.

The cell cycle and hypoxia meta-signatures were determined based on the respective Molecular Signatures Database (MsigDB^119^) and only correlated genes (R > 0.3) were considered. The relative expression of common signature genes between all samples was depicted in the expression heatmaps. For each cell cycle and hypoxia signature, a specific meta-module was defined, taking into account all genes that were common among the samples, and the average relative expression for each specific meta-module was calculated. These meta-modules were used to score the cells by the average relative expression of all genes in the meta-module, and cells were sorted according to these scores. The global score for each sample was calculated as the average of all cell cycle and hypoxia meta-modules expression. Meta-modules were also defined for the G1/S and G2/M phases of the cell cycle, which enabled cells to be classified as cycling (mean relative expression ≥ 0.1 and qval < 0.05) and noncycling (mean relative expression < 0.1 and qval > 0.05). For each cell, the mean relative expression of unique tumor subtype genes was calculated and used to create a score for each respective subtype. The minimum and maximum score values were determined and only cell scores above the threshold (qval > 0.001) were used to generate the tumor subtype heatmaps. Single cell signature scores for cellular phenotypic states and meta-modules *(MES, AC, NPC* and *OPC-like)* were implemented as described by Neftel *et al*., 2019^58^. TCGA subtypes of single cells were assessed based on signatures described in Wang *et al*., 2017^5^. Due to the limitations of Drop-seq data the signature scores for TCGA subtypes were determined according to scripts from Neftel *et al*., 2019^58^.

### Western Blot

Protein extraction was performed using minimal amounts of RIPA buffer (Thermo Fisher Scientific, Cat No. 89901) containing 1x protease inhibitor (Merck, cOmplete^®^ protease inhibitor cocktail) and on ice incubation for 15min followed by brief sonication and a centrifugation step (13.000 x g, 5min, 4°C) to remove cellular debris. iProtein extracts were resolved in NuPage^TM^ 4-12 % BisTris Protein Gels (Cat No. NP0321BOX, Thermo Fisher Scientific, MA, US), and blotted onto an Invitrolon^™^ PVDF (Thermo Fisher Scientific, Cat No. LC2005) or a Nitrocellulose membrane (Lifetech, Cat No. IB23001) according to standard protocols. After incubation with 5% nonfat milk in TBST (10 mM Tris, pH 8.0, 150 mM NaCl, 0.5% Tween 20) for 60 min, the membrane was rinsed with TBST and incubated with primary antibodies (**Table S7**). Membranes were washed three times for 10 min and incubated with horseradish peroxidase (HRP)-coupled secondary antibodies (Jackson ImmunoResearch) for 1 h at RT. Blots were washed with TBST three times, once with TBS, developed with a chemiluminescent substrate (ThermoFisher) and imaged with the ImageQuant 350 scanning system (cooled-CCD camera, GE Healthcare).

### *Ex vivo* compound screening in 384-well plate format

PDOX tumors were dissociated with the MACS Neural Dissociation kit (Miltenyi Biotec) according to manufacturer’s instructions. Mouse cells were removed with Mouse Cell Depletion kit (Miltenyi Biotec). Tumor cells were seeded 1000 cells/well in organoid medium in 384-well plates (PrimeSurface^®^, S-Bio) and cultured for 72h to allow organoid formation. Organoids (n=4 per PDOX model per drug concentration, n = 2 per mice) were treated with the following compounds: Erlotinib (EGFR, SelleckChem), Gefitinib (EGFR, SelleckChem), AZD3759 (EGFR, SelleckChem), AG-490 (JAK2, EGFR, SelleckChem), Daphnetin (EGFR, PKA/C, SelleckChem), Palbociclib (CDK4/6, SelleckChem), Abemaciclib (CDK4/6, SelleckChem), TMZ (Sigma) and 1,2:5,6-Dianhydrogalactitol (DAG, VAL-083, Delmar) in a fourfold and seven-point serial dilution series ranging from 1 μM to 1 mM (DAG, TMZ) or 12 nM to 48 μM (remaining inhibitors). After 3 (TMZ, DEG) or 6 (remaining inhibitors) days of incubation at 37°C in a 5% CO2 humidified incubator for respective inhibitors, cell viability and cytotoxicity were measured with CellTiter-Glo^®^2.0 and CellTox^™^-Green assays (Promega) respectively according to the manufacturer’s instructions with a ClarioStar plate reader (BMG Labtech). The relative cell viability for each dose was obtained by normalization with untreated control (DAG) or dimethyl sulfoxide (DMSO, Sigma, remaining compounds) per each plate or condition. Dose response curves (DRCs) were fitted using GraphPad Prism 8: best-fit lines and the resulting IC_50_ values were calculated using log[inhibitor] versus normalized response – variable slope (four parameters). The area under the curve (AUC) for each DRC was calculated using GraphPad Prism 8. For LIVE/DEAD double labeling, organoids were incubated with 2 mM Calcein-AM and 4 mM Ethidium homodimer-1 (LIVE/DEAD assay kit, Molecular Probes) for up to 6h. Imaging of viable (green) and dead (red) cells was done using LSM510 or LSM880 Confocal Laser microscopes (Zeiss).

### Cell printing and high through-put drug screening procedure

PDOX T434 tumors were dissociated with the MACS Neural Dissociation kit (Miltenyi Biotec) according to manufacturer’s instructions. Mouse cells were removed with Mouse Cell Depletion kit (Miltenyi Biotec). Tumor cells were mixed with 1% alginate (ratio of 1:1) and printed on 384-pillar array (1000 cells with 250nl) by ASFA Spotter ST (Medical & Bio Device, Suwon-si, South Korea). The pillars were washed by carefully combining the cell-pillar plates with 384-well plates containing 40 μL of cell culture medium (DMEM (Biowest), 10% FBS, 1% Pen-Strep, 4x NEAA (Lonza), 1% Ultraglutamine (Lonza)) in each well, and incubation for 30 min at 37°C. The pillar plates were then combined with 384-well assay plates containing cell culture medium and incubated for 3 days at 37°C and 5% CO2 atmosphere. The pillar plates were then transferred to compound plates where the cells immobilized in alginate were exposed for 7 days to 42 FDA-approved drugs, in a fourfold and seven-point serial dilution series from 7.3 nM to 30 μM in duplicates. To determine end-point cell viability, the cells were stained using Calcein AM live cell staining and the images were acquired using High Content imaging instrument (CV8000, (Yokogawa, Tokyo, Japan). Cell viability was calculated based on Calcein AM fluorescence. The relative cell viability for each dose was obtained by normalization with dimethyl sulfoxide (DMSO) per each plate. Dose Response Curves (DRC) were fitted using GraphPad Prism 8 (GraphPad). The AUC for each DRC was calculated using GraphPad Prism 8. Statistical differences between genetically defined groups were performed with unpaired 2-tailed t-test.

### *In vivo* tumor treatment

T16 GBM organoids were orthotopically implanted into the right frontal lobe of Swiss nude mice. Animals were monitored daily and the following criteria were evaluated: (1) loss of >10% of body weight, (2) exhibition of strong neurological signs (3) increased lordosis or (4) swollen belly. Tumor growth was monitored by MRI (T1- and T2-weighted MRI protocol; 3T MRI system, MR Solutions). 35 days postimplantation most mice had visible tumors and were randomized into 4 treatment groups (7 mice per treatment group, 6 mice per control group): Control, Bevacizumab (Avastin) treatment, Dianhydrogalactitol (DAG, VAL-083) treatment and combined Bevacizumab + DAG treatment. Drug concentrations and treatment schedule were as follows: Bevacizumab - 20mg/kg, 1x week, DAG - 3.5mg/kg, 3x week. Control animals received saline (NaCl 0.9%) 4 x week. Compounds and saline were delivered by intraperitoneal injections. Bevacizumab and DAG injections were performed on different days. 49 days after implantation MRI T2 was applied to monitor tumor progression. T1 with contrast agent was applied to several mice to evaluate the response of tumor to Bevacizumab. 56 days after implantation one mouse in control group showed neurological symptoms and was euthanized directly after MRI. T2 and T1 + contrast MRI was applied to all mice. Remaining mice were euthanized the following day before mice developed symptoms and brains extracted. Tumor volume (mm^3^) was measured in ImageJ as the sum of area obtained by tumor delineation in each slice and multiplying by slice thickness (1mm). Growth rate (GR) was calculated using the TV measurement as GR = 100 * log (TVf/TV0) / (tf-t0), where TVf and TV0 are the tumor volumes at the late (day 56) and early (day 35 or day 49) time points respectively, and tf-t0 is the difference in days between the time points. Tumor volumes are expressed in mm3 and GR in ‘% per day^120^.

### Statistical tests

Different statistical approaches have been applied based on the data type and measurements across the manuscript. Statistical tests are described in each paragraph above corresponding to the associated experimental procedures. If not specified above, significant differences were calculated with the Student’s t-test.

### Data Availability

Molecular data are available in the Gene Expression Omnibus repository (https://www.ncbi.nlm.nih.gov/geo/) under accession numbers as follows: (1) array-CGH: GSE137959; (2) DNA methylation: GSE137845; (30 gene expression: GSE134470; (4) scRNA-seq:GSE128195. targeted DNA sequencing is available in the Sequence Read Archive (https://www.ncbi.nlm.nih.gov/sra) under accession number SUB7313530. Remaining datasets supporting the findings are available from the corresponding author on reasonable request. PDOX models are available from the corresponding author or via EurOPDX consortium (https://www.europdx.eu/) and PDXFinder (https://www.pdxfinder.org/^121^; new release May 2020).

## ACKNOWLEDGEMENTS

We are grateful to the Clinical and Epidemiological Investigation Center (CIEC) of the LIH for support in tumor collection, and to Jean-Jacques Gerardy and Nathalie Nicot for technical assistance. We thank the Helmholtz Zentrum München (Research Unit of Molecular Epidemiology/Institute of Epidemiology, German Research Center for Environmental Health, Neuherberg, Germany) for processing the EPIC DNA methylation arrays.

We acknowledge the financial support by the Luxembourg Institute of Health, Télévie-FNRS (Grants n°7.4632.17, 4615.18), Fondation Cancer Luxembourg (Pan-RTK Targeting), the Luxembourg National Research Fund (FNR; CORE Junior C17/BM/11664971/DEMICS) and the GLIOTRAIN ITN funded by the European Union’s Horizon 2020 research and innovation programme under the Marie Skłodowska-Curie grant agreement No 766069 (The material presented and views expressed here are the responsibility of the author(s) only. The EU Commission takes no responsibility for any use made of the information set out). MM would like to thank the Luxembourg National Research Fund (FNR) for the support (FNR PEARL P16/BM/11192868 grant).

## AUTHOR CONTRIBUTION

Conceptualization: A.G., A-C.H., A.O., R.B., S.P.N; Methodology: A.G., A-C.H., A.O., D.S., Al.S., S.P.N Investigation: A.G., A-C.H., A.O., D.S., Y.A.Y, Vi.B., Va.B., E.K., S.B., O.K., Al.M., A. DF., L.R., S.P., Al.M., M.M., Formal analysis: A.G., A-C.H., A.O., D.S., Y.A.Y., V. N., M.W., Ar.M., T.K., P.V.N., F.A., A.DF., B.F., S.P., T.A., Ar.M., P.M., H.M., T.M.M., H.N., Y-J.K., W.J., B. K., G.T., L.F.S., Resources: Al.M., C.H-M., An.S., D.B., H.M., Al.S., F.H., R.B., S.P.N, Supervision: A.G., C.H-M., B.K., L.F.S., M.M., Al.S., R.B., S.P.N., Writing - Original Draft: A.G., A-C.H., S.P.N., Writing - Review & Editing: all authors

## DECLARATION OF POTENTIAL CONFLICT OF INTEREST

D. Brown is CSO at Delmar Pharmaceuticals, A. Steino was preclinical study manager at Delmar Pharmaceuticals. A Golebiewska and S.P. Niclou report receiving a commercial research grant from Delmar Pharmaceuticals. No potential conflict of interest was disclosed by other authors.

