## Supplementary Figures and Tables for "Primary and recurrent glioma patient-derived orthotopic xenografts (PDOX) represent relevant patient avatars for precision medicine"

### FIGURE S1

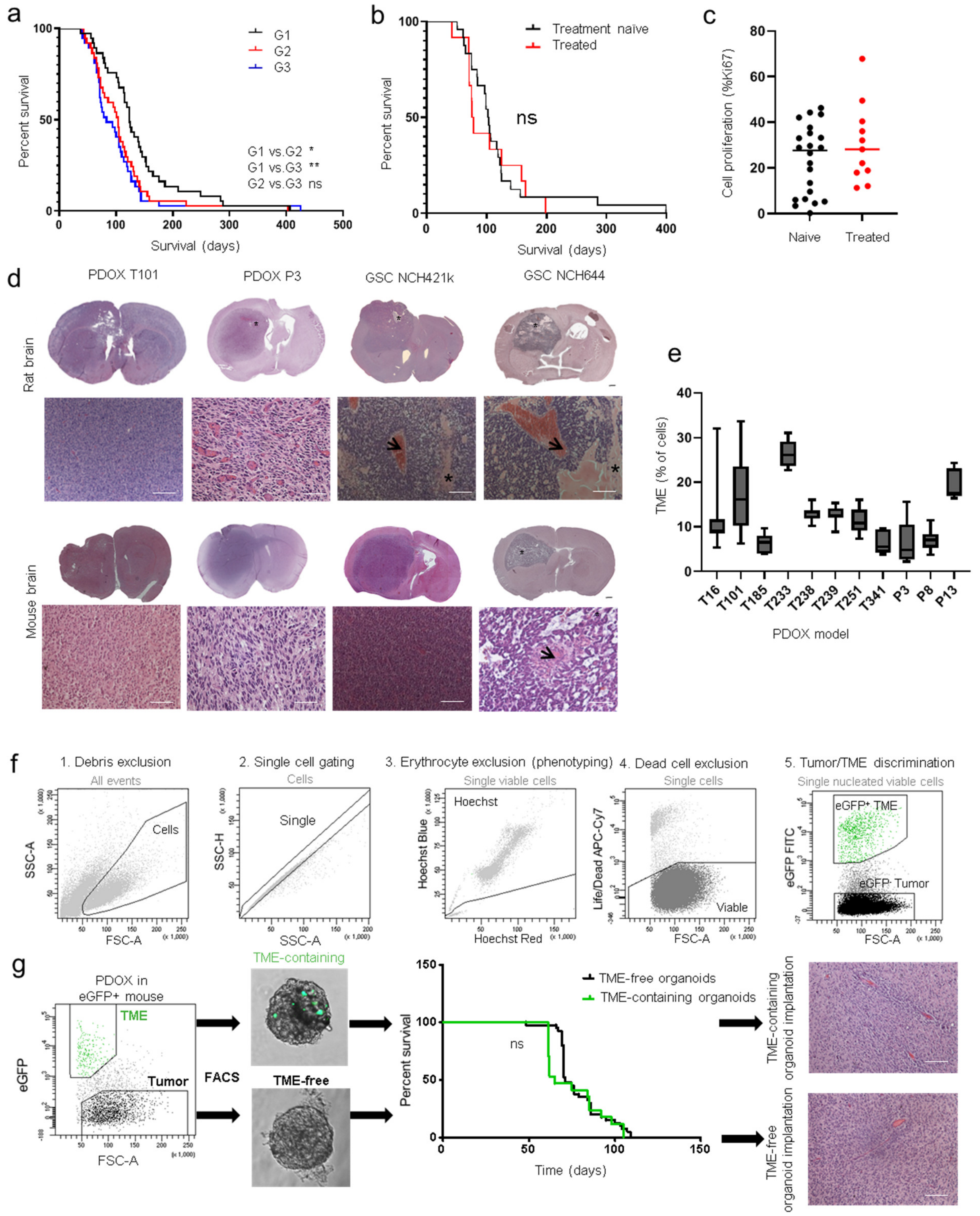


**Figure S1. Characterization of glioma PDOX models. a** Kaplan-Meier survival curves of PDOXs at generation 1 (G1), 2 (G2) and 3 (G3) for those models where G ≥ 3 was reached (n=37). Mean survival of each model per generation was plotted in each group. See **Table S1** for details; (* p-value <0.05, ** p-value <0.01, Wilcoxon signed-rank test). **b** Kaplan-Meier survival curves of PDOXs separated into treatment-naïve and treated gliomas. ns = not significant (Log-rank and Wilcoxon signed-rank tests). **c** Comparison of cell proliferation in PDOXs (mean %Ki67 per model) derived from treatment-naïve and treated patient tumors. No statistically significant difference was observed (unpaired two-tailed t-test). **d** Additional examples showing variance of histopathological features in rats and mice. Comparison of PDOXs (T101, P3) and GSC line-derived xenografts (NCH421k, NCH644) with strong invasive (T101), towards intermediate (P3, NCH421k) and angiogenic (NCH644) features (arrows = microvascular proliferation, stars = pseudopalisading necrosis, black bar = 1 mm, white bar = 100 µm). **e** Quantification of mouse derived tumor microenvironment (TME) content in the tumor core of PDOX models. Mouse cells were recognized by flow cytometry as hCD90 negative viable cells. **f** Gating strategy for flow cytometry**.** Example is shown for PDOX P3 in eGFP$^+$ NOD/SCID mice (1) Cells were distinguished from debris based on the Forward Scatter (FSC) and Side Scatter (SSC). (2) Cell aggregates were gated out based on their properties displayed on the SSC area (SSC-A) versus height (SSC-H) dot plot. (3) For multicolor phenotyping, erythrocytes were excluded on the 'Hoechst Red'/'Hoechst Blue' dot plot in the linear scale. Hoechst staining was omitted for sorting due to increased toxicity (4) Dead cells were recognized by their strong positivity for the dead cell marker (5) In PDOXs, human tumor cells were recognized as eGFP negative, compared to the eGFP positive mouse non-malignant cells (tumor microenvironment = TME). **g** TME-free organoids were derived from FACS-sorted PDOX T16 eGFP- tumor cells only. As control, TME-containing organoids were generated from simultaneous FACS of eGFP- tumor and eGFP+ TME cells. Kaplan-Meier survival curves represent survival of mice across 5 generations per group (G1-G5, total n = 40 for TME-containing and n =12 for TME-free passaging). No statistical difference was observed (Log-rank and Wilcoxon signed-rank tests). Representative histology of PDOXs obtained by TME-containing and TME-free passaging (generation 4) shows no change in tumor phenotype.

### FIGURE S2

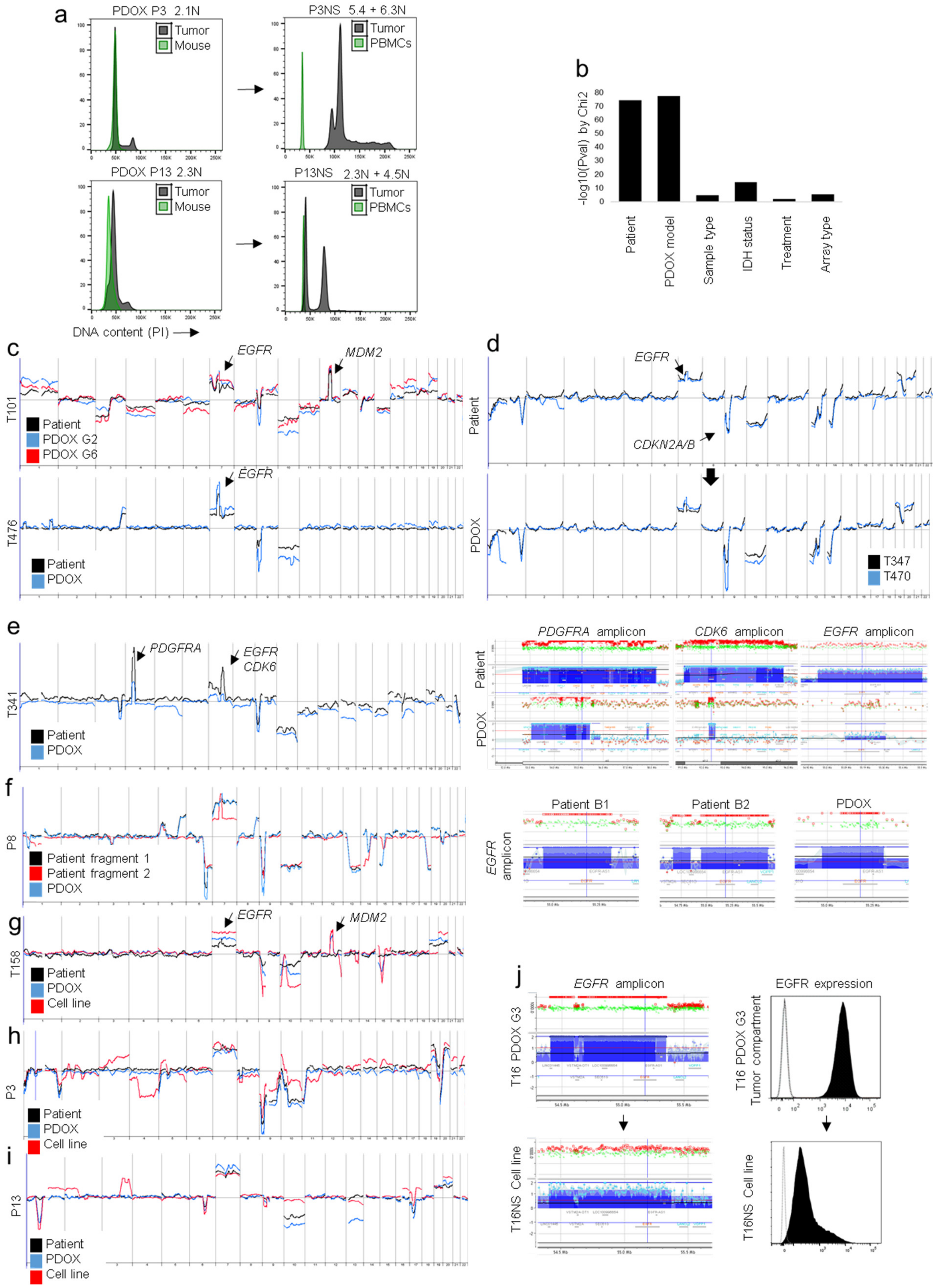


**Figure S2. Genetic aberrations in glioma preclinical models. a** Ploidy analysis of tumor cells in P3 and P13 glioma PDOXs and PDOX-derived GSC cultures (P3NS, P13NS). Mouse cells were used as diploid control (2N) in PDOX samples, PBMCs were used as diploid control (2N) for *in vitro* cultures. Examples are shown for cultures in passage < 10. See more examples in **Table S1**. Note strong aneuploidization of P3 and P13 GSC lines upon *in vitro* cultures (P3NS, P13NS). **b** Statistical analysis of array-CGH data. $Chi^2$ test for independence reveals limited impact of sample type, treatment and array type on genetic profiles. Individual tumor genetic aberrations and *IDH1* mutation status are main sources of variation. **c** Examples of array-CGH profiles of GBM patient tumors and corresponding PDOX models are shown for T101 and T476. Genetic aberrations were recapitulated over serial transplantations. See **Table S2** for detailed description. **d** Array-CGH profiles of longitudinal samples (T347-T470) of GBM patient LIH0347 showing the same genetic aberrations upon recurrence. These profiles are largely recapitulated in PDOX models, only an additional 1p31.1-p11.2 loss was detected in PDOX T470. **e** Array-CGH profiles of GBM T341 patient tumor and corresponding PDOX. PDOX model was derived from an additional *MDM4/CDK6*-amplified clone with different chromosomal breakpoints. Right panels show presence of different amplicons in the patient tumor and corresponding PDOX. **f** Array-CGH profiles of GBM P8 patient tumor fragments and corresponding PDOX. Analysis of 2 tumor fragments revealed intra-tumoral genetic heterogeneity and different *EGFR* amplicon. **g-i** Array-CGH profiles of GBM patient tumors, corresponding PDOXs and *in vitro* GSC lines for T158 (g), P3 (h), P13 (i). Additional aberrations occurred upon *in vitro* passaging. Note that PDOX T158 arose from an additional *MDM2*-amplified, *EGFR*-non amplified clone, not detected in the patient sample. **j** *In vitro* passaging of T16 tumor cells as GSC line (T16NS) led to loss of *EGFR* amplicon (array-CGH, left panel) and decreased EGFR expression (flow cytometry, right panel).

**FIGURE S3**

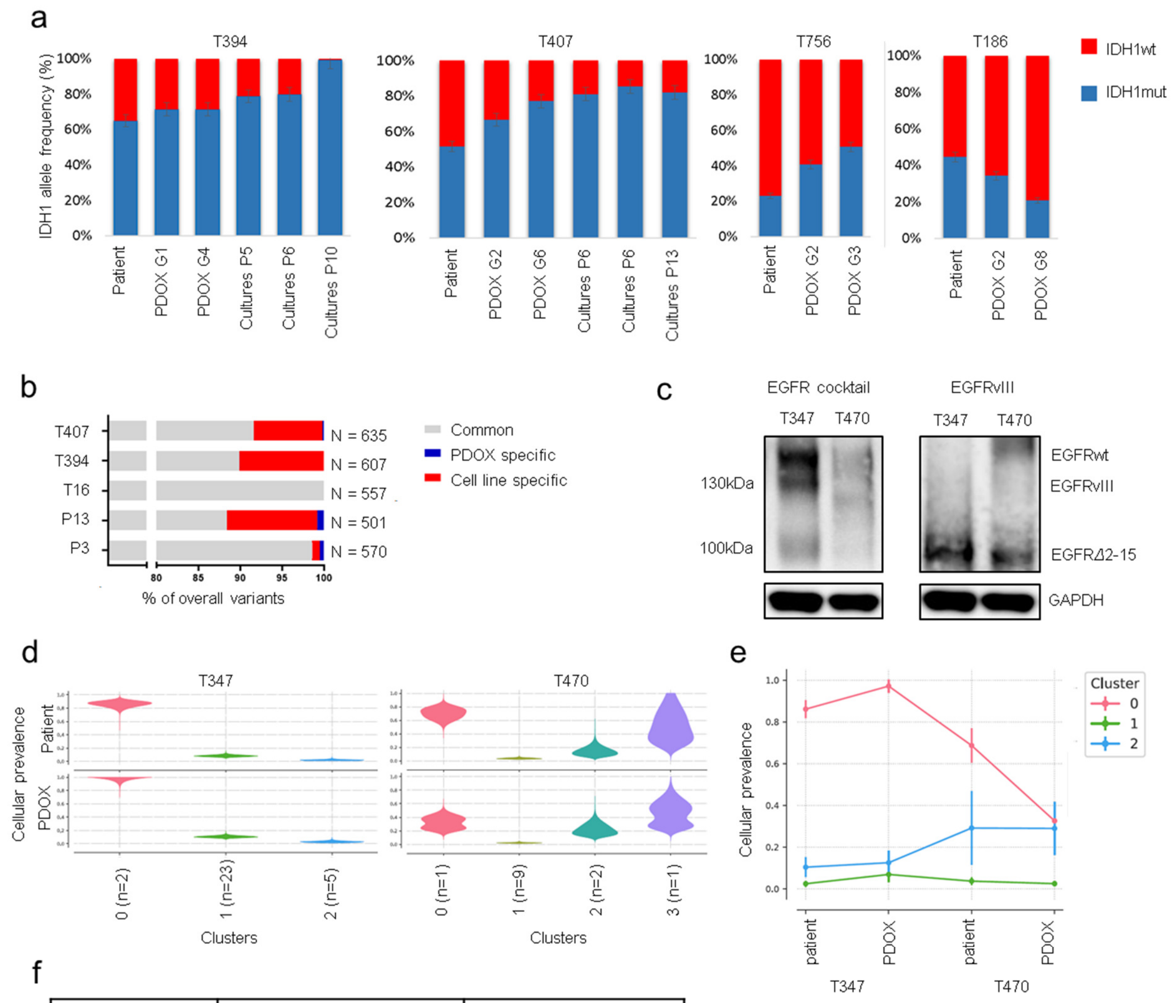


| Variant EGFR | chr7:55168634 G>C | | chr7: 55154017 C->T | |
|---|---|---|---|---|
| Sample | Read depth | Alt reads | Read depth | Alt reads |
| T192_patient | 983 | 3 | 5966 | 330 |
| T192_PDOX | 1169 | 988 | 6287 | 375 |
| T233_patient | 401 | 0 | 705 | 0 |
| T233_PDOX | 625 | 0 | 1346 | 0 |
| T251_patient | 2245 | 0 | 4422 | 2 |
| T251_PDOX | 1826 | 0 | 1865 | 1 |

**Figure S3. Recapitulation of genetic heterogeneity in glioma PDOXs. a** Digital PCR-based analysis of *IDH1 wild-type* and *R132H* fractions in IDH1mut gliomas and corresponding preclinical models. **b** Recapitulation of overall variants detected by targeted sequencing. Cell lines were compared to respective patient samples. **c** Western blots against EGFR (antibody cocktail recognizing wt and structural variants) and EGFRvIII proteins in PDOXs T347 and T470 (patient LIH0347) show wildtype EGFR protein as well as the structural variant EGFRΔ2-15, which is recognized by EGFRvIII antibody, despite decreased molecular weight. **d** Cellular prevalence estimates from PyClone representing clonal subpopulations detected in patient tumors and respective PDOXs. Examples shown for longitudinal samples (T347, T470) of patient LIH0347. Each cluster of mutations was computationally inferred to reflect a subclone. Number of genetic variants contributing to each clone is depicted. **e** Cellular prevalence estimates from PyClone representing clonal subpopulations detected in longitudinal samples of patient LIH0347 and the respective PDOXs. Each line represents a cluster of mutations computationally inferred to reflect a subclone. Only genetic variants detected in all samples were considered for analysis. **f** Evolutionary dynamics of *EGFR* genetic variants. Targeted DNA sequencing revealed longitudinal evolution of *EGFR* genetic variants in LIH0192 patient tumors and PDOXs derived thereof. A specific subclonal variant was present only in T192 patient tumor and was enriched in the respective PDOX (55168634 G->C). Another genetic variant present in T192 patient tumor and PDOXs at the subclonal level was selected out during disease progression (chr7:55154017 C>T). Comparison of overall read depths suggests that these variants are discordantly inherited upon tumor recurrence in patients and PDOX derivation, supposedly via unequal distribution of extrachromosomal DNA.

### FIGURE S4

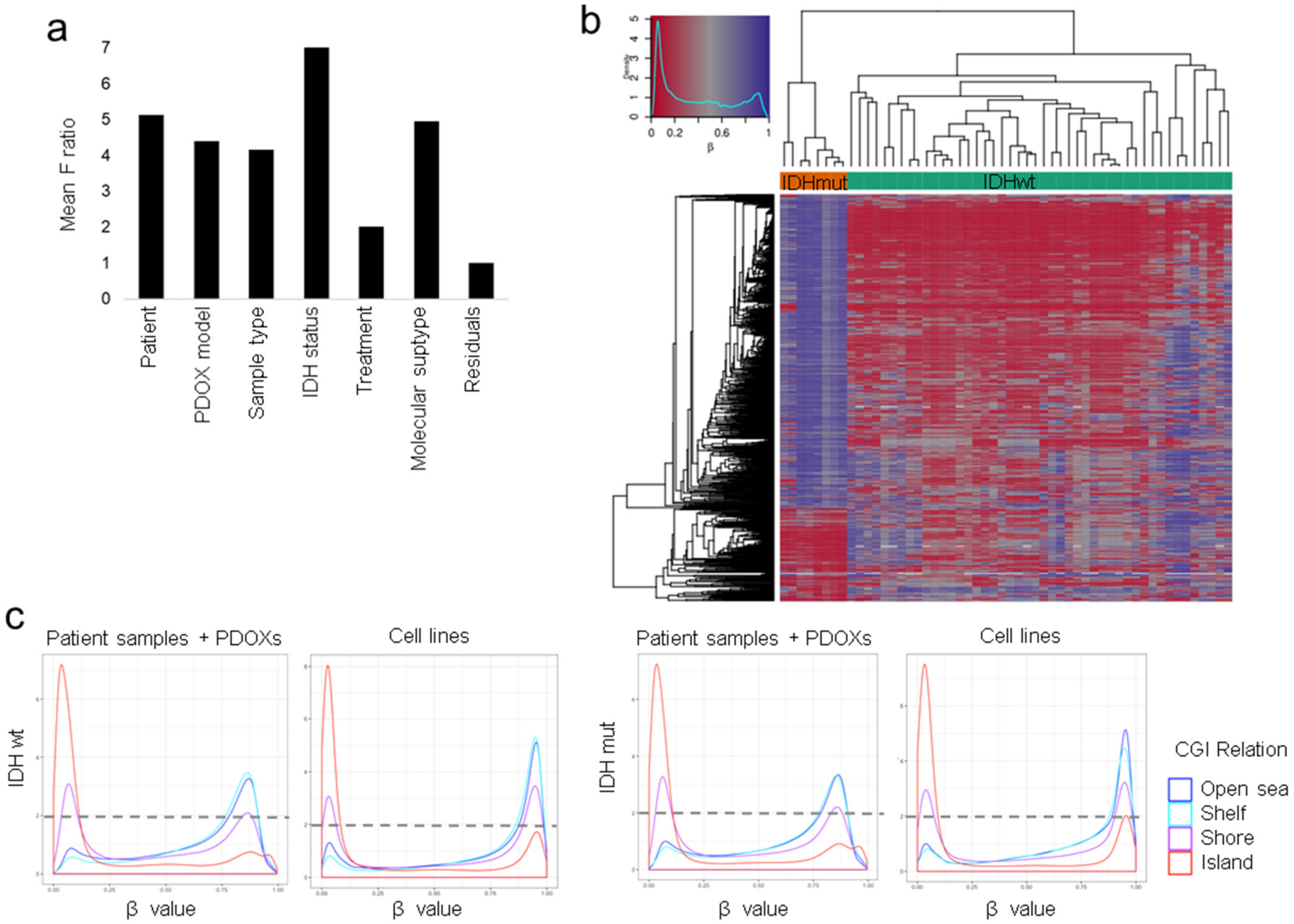


**Figure S4 Recapitulation of DNA methylation profiles in glioma PDOXs**. **a** Single factor ANOVA based on global beta-value distributions reveals IDH mutation status as a main source of variation in the DNA methylation cohort. **b** Heatmap representing 1000 most variable features in DNA methylation shows genomic loci differentially methylated between IDH1mut versus IDHwt patients and preclinical models. **c** beta-value distributions are very similar between IDH1mut and IDHwt tumors (patient samples and PDOX models), in accordance with the G-CIMP low status of the IDH1mut gliomas. IDHwt and IDH1mut GSC lines increase DNA methylation at numerous sites corresponding to open seas, shelfs, and shores.

### FIGURE S5

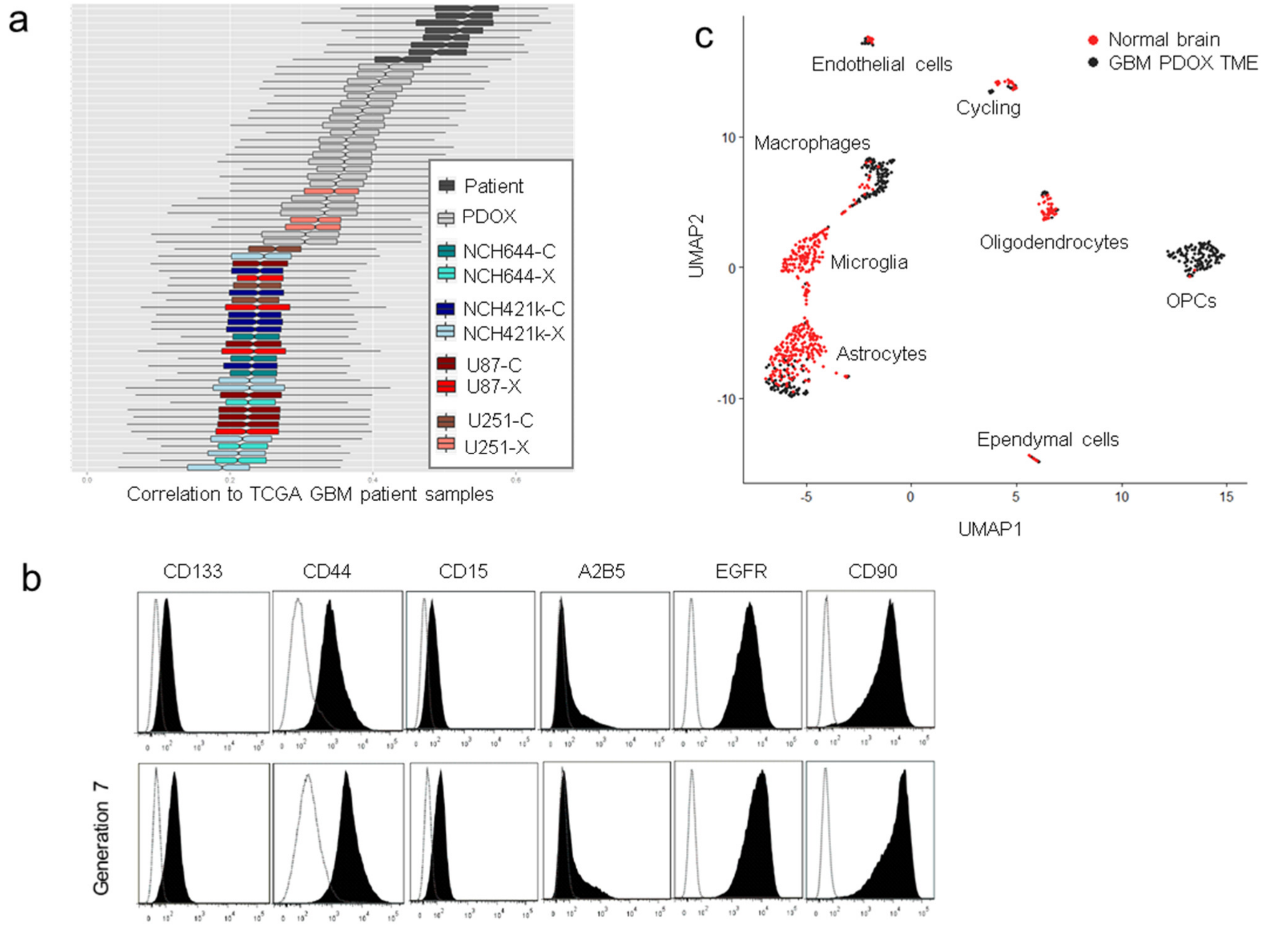


**Figure S5. Gene expression profiles in glioma preclinical models. a** Correlation of gene expression profiles to TCGA GBM patients shows close resemblance of PDOXs to patient tumors at the transcriptomic level. **b** Stem-cell associated marker expression profiles were interrogated by flow cytometry in tumor cells of PDOXs over serial transplantation. Example shown for PDOX T101. **c** Single cell RNA-Seq of mouse brain showing overall gene expression relationship between cells of normal brain (red) and upon GBM implantation (shown for PDOX P8) (black). Identified TME subpopulations are depicted.

**FIGURE S6**

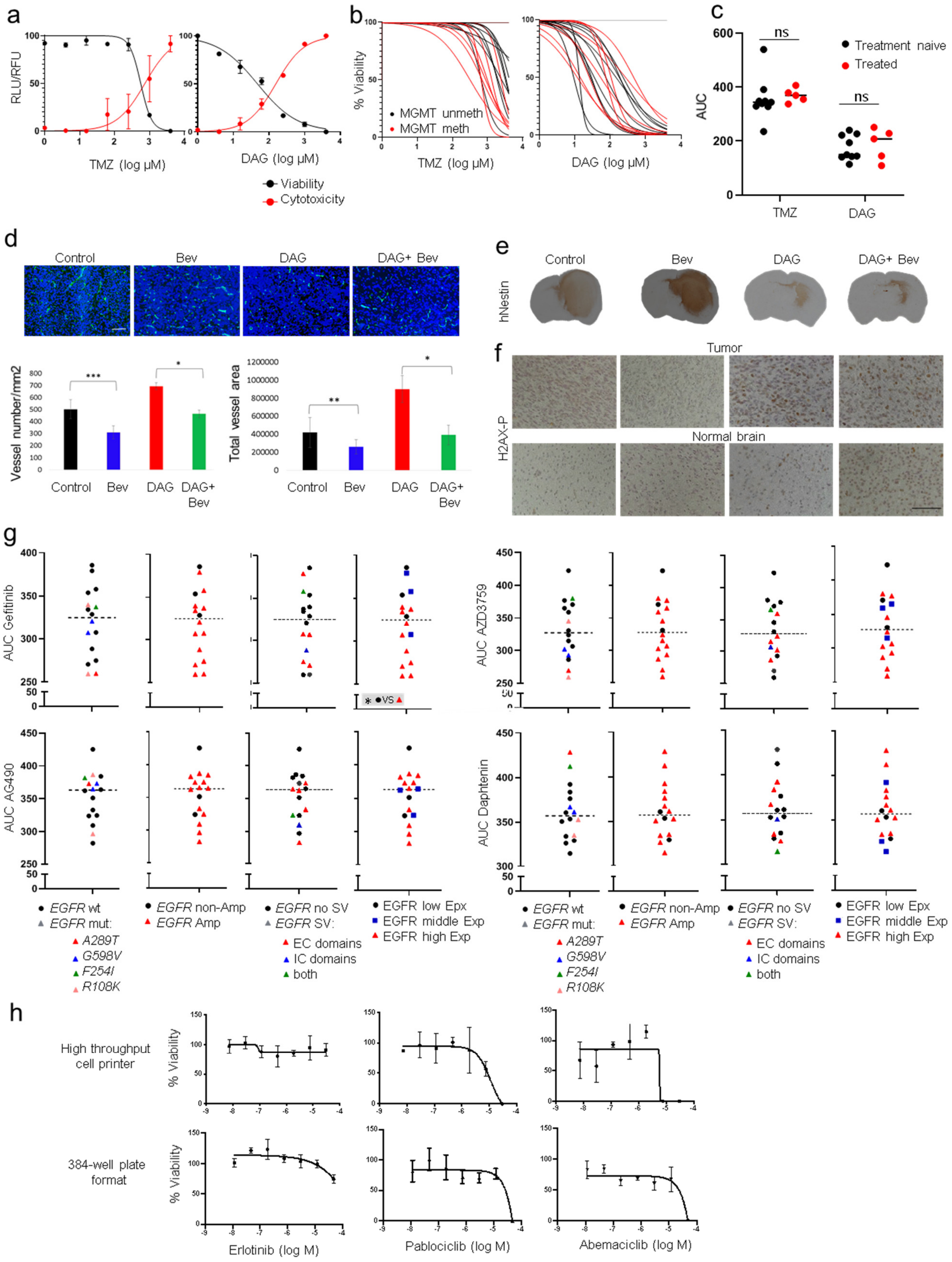


**Figure S6. Drug testing regimens *ex vivo* and *in vivo*. a** Quantification of cell viability and toxicity displayed as normalized and log transformed RLU/RFL. Examples are shown for TMZ and DAG response in P3-derived organoids. **b** Response curves (non-linear fit, n = 3) of individual PDOXs to TMZ and DAG treatment displayed as %-Viability normalised to untretated control. *MGMT* promoter unmethylated and methylated tumors are shown in black and red respectively. **c** Mean AUC upon exposure to TMZ and DAG in PDOX models derived from treatment-naïve and treated tumors. ns = not significant (unpaired t-test). **d** Blood vessels *in vivo* were visualized with mouse specific anti-CD31 staining (in green). Tumor was defined as nuclei dense area (nuclei in blue = DAPI). Representative pictures are shown for each experimental group (Scale bar = 100µm). Quantification of vessel number per $mm^2$ upon treatment and area covered by vessels confirmed normalization of the tumor vasculature in Bevacizumab (Bev) treated mice (Mean±SD, *$p_{value}$<0.05, ***$p_{value}$ <0.01, ****$p_{value}$ <0.001, n = 5-6 for Control and Bev groups, n = 2 for DAG treated groups, 3-5 pictures were taken per each tumor, unpaired t-test). The statistical analysis between Control and DAG treated groups is not shown due to major differences in tumor volumes, leading to lower aberrations in tumor vasculature of the small DAG treated tumors. **e** Representative sections of PDOX stained against human-specific Nestin (n = 3). **f** IHC for H2AX-P in PDOX sections (brown nuclei = H2AX-P). Counterstaining for nuclei with hematoxyline. Induction of H2AX-P was observed in DAG treated tumors. Minor induction of H2AX-P was also observed in a subpopulation of normal brain cells. Scale bar = 100 µm. **g** Quantification of AUC upon exposure to EGFR inhibitors: Gefitinib, AG490, AZD3759 and Daphtenin (*$p_{value}$ < 0.05, unpaired t-test); *wt = wildtype, mut = mutated, Amp = amplified, SV = structural variant, exp = protein expression.* **h** Response curves (non-linear fit, n = 3 for plate formar, n = 2 for cell printer) of PDOX T434 to EGFR and CDK4/6 inhibitors displayed as %-Viability. Similar treatment responses were observed with the two protocols applied.

### SUPPLEMENTARY TABLES

**Table S1.** **Clinical data corresponding to patients of which PDOX models were derived.** Clinically relevant patient information is displayed in columnt C-J, such as age at tumor collection, sex, tumor localization based on MRI, treatment information, histological and molecular diagnosis (https://www.molecularneuropathology.org/mnp) of patient tumors. IDH status as assessed by NGS and MGMT promoter methylation status from Illumina Infinium HumanMethylation BeadChips.
Further, PDOX model relevant info is presented in column K to O, such as implanted tissue type, generations reached, stable survival time, cell proliferation index and ploidy.
If available, PDOX-derived GSC lines are specified in columns P-Q; NA – Not available.

| PDOX model | Patient | Age at collection | Sex | Tumor localization | Treatment received prior PDOX | Histological diagnosis | Molecular diagnosis | IDH1 status (patient) | MGMT methylation (patient) | Implantation | PDOX generation reached | PDOX survival time (days) | Proliferation index in PDOX (%Ki67 +/- SD) | Ploidy PDOX | PDOX-derived cell line | Ploidy cell line |
|---|---|---|---|---|---|---|---|---|---|---|---|---|---|---|---|---|
| P3 | BER0003 | 64 | Male | NA | NA | GBM Grade IV | NA | WT | methylated | spheroids | >G10 | 42.5 +/- 3.6 | 54.7 +/- 3.7 | 2.1N | P3NS | 5.4N + 6.3N |
| P8 | BER0008 | 64 | Female | NA | NA | GBM Grade IV | GBM, IDHwt, RTK I | WT | methylated | spheroids | >G10 | 57.4 +/- 5 | 38.3 +/- 10.8 | 2.2N | | |
| P13 | BER0013 | NA | Female | NA | NA | GBM Grade IV | NA | WT | methylated | spheroids | >G10 | 36.5 +/- 4 | 42.9 +/- 10.8 | 2.3N | P13NS | 2.3N + 4.5N |
| T16 | LIH0016 | 52 | Female | Fronto-parietal right | TMZ | GBM Grade IV | GBM, IDHwt, mesenchymal | WT | unmethylated | spheroids | >G10 | 75 +/- 6.5 | 18.9 +/- 5.8 | (2.1N) + 3.4N | T16NS | 3.8N |
| T101 | LIH0101 | 60 | Male | NA | NA | GBM Grade IV | NA | WT | methylated | spheroids | G7 | 101 +/- 11 | 13.4 +/- 4.8 | (2N) + 4.6N | | |
| T158 | LIH0158 | 70 | Female | Frontal right | no prior treatment | GBM Grade IV | GBM, IDHwt, mesenchymal | WT | methylated | spheroids | G5 | 77+/- 18 | 35.2+/-3 | 2.3N | T158NS | 2.4N |
| T185 | LIH0185 | 76 | Female | NA | no prior treatment | GBM Grade IV | GBM, IDHwt, RTK II | WT | methylated | spheroids | G6 | 89 +/- 10 | 19.4 +/- 6.9 | 2.2N | | |
| T186 | LIH0186 | 34 | Male | NA | no prior treatment | Anaplastic Oligodendroglioma Grade III | Glioma, IDHm, high grade astrocytoma | Mutant R132H | methylated | fresh spheroids/tissue | G9 | 159 +/- 43 | 33+/-4.8 | 2.1-2.4N | | |
| T188 | LIH0188 | 68 | Male | NA | no prior treatment | GBM Grade IV | GBM, IDHwt, RTK II | WT | unmethylated | spheroids | G5 | 64 +/- 10 | 37.1+/- 2.9 | 2.3N | | |
| T192 | LIH0192 | 42 | Female | Frontal left | no prior treatment | GBM Grade IV | GBM, IDHwt, mesenchymal | WT | unmethylated | spheroids | G6 | 66 +/- 6 | 26.5 +/-8 | 2.4N | | |
| T226 | LIH0188 | 69 | Male | NA | radiotherapy +TMZ | GBM Grade IV | NA,low tumor content | WT | NA | spheroids | G3 | 142 +/- 23 | NA | NA | T226NS | 2.0N + 3.7N |
| T233 | LIH0192 | 42 | Female | Frontal left | radiotherapy + TMZ | GBM Grade IV | GBM, IDHwt, mesenchymal | WT | unmethylated | spheroids | G4 | 121 +/- 9 | 11,3 | 2.3N | | |
| T238 | LIH0238 | 42 | Male | NA | no prior treatment | GBM Grade IV | GBM, IDHwt, inflamatory tissue | WT | methylated | spheroids | G3 | 119 +/- 32 | 22.1 | (2.1N) + 3.7N | | |
| T239 | LIH0239 | 80 | Male | NA | no prior treatment | GBM Grade IV | GBM, IDHwt, mesenchymal | WT | methylated | spheroids | G3 | 124 +/- 25 | 4.5+/-4.9 | 2.3N | | |
| T251 | LIH0192 | 43 | Female | Frontal left | radiotherapy +TMZ | GBM Grade IV | GBM, IDHwt, RTK II /mesenchymal | WT | unmethylated | spheroids | G6 | 72 +/- 11 | 36.1 +/- 7.7 | 2.3N | | |
| T281 | LIH0281 | 51 | Female | Parietal Left | no prior treatment | GBM Grade IV | GBM, IDHwt, mesenchymal | WT | methylated | spheroids | G3 | 117 +/- 13 | 5.3 +/-0.41 | 2.0N + 3.5N | | |
| T304 | LIH0304 | 53 | Male | Frontal left | no prior treatment | GBM Grade IV | GBM, IDHwt, RTK II /I | WT | unmethylated | spheroids | G3 | 84 +/- 15 | 6.01 | 2.2N + 3.2N +4.4N | | |
| T331 | LIH0331 | 84 | Male | Fronto-temporo-parietal left | no prior treatment | GBM Grade IV | GBM, IDHwt, mesenchymal | WT | methylated | spheroids | G4 | 116 +/- 5 | 33.2 +/- 2.73 | 2.0N | | |
| T341 | LIH0337 | 75 | Female | Temporal right | no prior treatment | GBM Grade IV | GBM, IDHwt, RTK I | WT | methylated | spheroids | G4 | 50 +/- 5 | 37.7 +/- 6.1 | 2.1N + 3.65N | | |
| T347 | LIH0347 | 41 | Male | Frontal left | no prior treatment | GBM Grade IV | GBM, IDHwt, RTK II | WT | unmethylated | spheroids | G3 | 119 +/- 21 | 42.1+/-1.9 | 2.2N | | |
| T356 | LIH0281 | 52 | Female | Parietal Left, para-sagittal | radiotherapy +TMZ | GBM Grade IV | GBM, IDHwt, mesenchymal | WT | methylated (low tumor content) | spheroids | G4 | 162 +/- 45 | 28.17 +/- 4.49 | 2.1N | | |
| T361 | LIH0361 | 69 | Female | Temporal left, para-brainstem | no prior treatment | GBM Grade IV | GBM, IDHwt, RTK II | WT | methylated | spheroids | G5 | 75.5 +/- 8 | 46.3+/-1.6 | 2.3N | | |
| T363 | LIH0363 | 84 | Female | Occipital lobe right | no prior treatment | GBM Grade IV | NA | WT | NA | spheroids | G3 | 138 +/- 2 | 6.36 | 2.1N | | |
| T367 | LIH0367 | 69 | Male | Temporal left | no prior treatment | GBM Grade IV | GBM, IDHwt, RTK II | WT | methylated | spheroids | G2 | 140 +/- 0 | 29.7+/-3.1 | 2.3N | | |
| T384 | LIH0384 | 50 | Female | Temporo-Occipital left | no prior treatment | GBM Grade IV | GBM, IDHwt, mesenchymal | WT | methylated | spheroids | G2 | 225 +/- 51 | NA | NA | T384NS | 3.65N |
| T386 | LIH0386 | 51 | Male | Frontal left | no prior treatment | GBM Grade IV | GBM, IDHwt, RTK I | WT | unmethylated | spheroids | G4 | 116 +/- 21 | 28.9+/-0.72 | 2.1N | | |
| T394 | LIH0394 | 45 | Female | Fronto-basal right | radiotherapy | GBM Grade IV | Glioma, IDHm, high grade astrocytoma | Mutant R132H | methylated | fresh spheroids/tissue | G5 | 70 +/- 9 | 67.8+/-8.3 | 2.1-2.3N + 3.5N | T394NS | 3.1N |
| T407 | LIH0394 | 45 | Female | Fronto-basal right | radiotherapy + Avastin | GBM Grade IV | Glioma, IDHm, high grade astrocytoma | Mutant R132H | methylated | fresh spheroids/tissue | G4 | 68 +/- 4 | 49.5+/-6.8 | 2.1N + 3.9N | T407NS | 3.1N +3.5N |
| T434 | LIH0304 | 54 | Male | Frontal left | radiotherapy + TMZ | GBM Grade IV | GBM, IDHwt, RTK II | WT | unmethylated | spheroids | G6 | 44 +/- 3 | 40.4 +/- 11.43 | 2.0N + 2.3N +3.8N | | |
| T470 | LIH0347 | 42 | Male | Frontal left | radiotherapy + TMZ | GBM Grade IV | GBM, IDHwt, RTK II | WT | unmethylated | spheroids | G4 | 72 +/- 8 | 17.98 +/- 0.88 | 2.2N | | |
| T476 | LIH0476 | 75 | Male | Fronto-parietal left | no prior treatment | GBM Grade IV | GBM, IDHwt, RTK II/I | WT | unmethylated | spheroids | G4 | 63 +/- 6 | 44.3+/-3.8 | 2.3N | | |
| T515 | LIH0515 | 46 | Male | Frontal-right | no prior treatment | Anaplastic Oligodendroglioma Grade III | NA | Mutant R132H | NA | fresh spheroids/tissue | G3 | 413 +/- 16 | NA | NA | | |
| T591 | LIH0384 | 52 | Female | Temporo-Occipital right | radiotherapy +TMZ + Avastin | GBM Grade IV | GBM, IDHwt, RTK II /mesenchymal | WT | unmethylated | spheroids | G3 | 108 +/- 3 | 22.08+/-12.72 | 2.2N + 3.5N | | |
| T744 | LIH0744 | 69 | Male | Bifrontal | no prior treatment | GBM Grade IV | NA | WT | NA | spheroids | G3 | 103 +/- 4 | 28.9+/-23.5 | 2.2N | | |
| T756 | LIH0756 | 46 | Male | Multifocal left | radiotherapy | GBM Grade IV | Glioma, IDHm, high grade astrocytoma | Mutant R132H | methylated | fresh spheroids | G5 | 73 +/- 14 | 12.09 +/-1.62 | 2.2N + 2.9N + 3.9N | | |
| T772 | LIH0615 | 54 | Female | Fronto-temporal left | radiotherapy + TMZ | GBM Grade IV | GBM, IDHwt, RTK II | WT | methylated | spheroids | G4 | 106 +/- 6 | 32.1+/-1.2 | 2.1N | | |
| T784 | LIH0784 | 56 | Male | Frontal left | no prior treatment | GBM Grade IV | GBM, IDHwt, RTK I | WT | methylated | spheroids | G4 | 136 +/- 16 | 43.5+/-3.3 | 2.1N | | |
| T797 | LIH0797 | 55 | Female | Left occipital | no prior treatment | GBM Grade IV | GBM, IDHwt, RTK I | WT | unmethylated | spheroids | G3 | 126 +/- 9 | 9.65 +/-0.40 | 2.2N + 3.9N | | |
| T831 | LIH0831 | 51 | Female | Bifrontal | no prior treatment | GBM Grade IV | GBM, IDHwt, RTK II | WT | methylated | spheroids | G3 | 75 +/- 7 | 3.46 +/- 2.33 | NA | | |
| T832 | LIH0831 | 51 | Female | Bifrontal | no prior treatment | GBM Grade IV | GBM, IDHwt, RTK II | WT | methylated | spheroids | G2 | 123 +/- 2 | 0.25 +/- 0.04 | 2.1N | | |

**Table S2.** **Chromosomal aberrations of glioma patient samples and corresponding PDOX models and GSC lines.**

Modifications in genome structure as identified by array-CGH are shown for each patient tumor [++ = amplification (Log2 Ratio>2), + = gain (Log2 Ratio >0.25), - = loss (Log2 Ratio < -0.25), -- = deletion (Log2 Ratio< -1)]. If present, changes of genetic aberrations of patient profiles are shown for PDOX models and cell lines in column c and d respectively . Genetic aberrations determined only from EPIC methylation arrays are highlighted with red background.

| PDOX model | Patient tumor | PDOX specific changes vs. patient | Cell line specific changes vs. PDOX |
|---|---|---|---|
| P3 | +[Chr7, 19p, 20q], -[1p36-1p34.1, 1q21.1-q44, -5p15.33-31, Chr9, Chr10, 11p15-14, 20p] --CDKN2A/B | | +(3q, Chr21, Chr22), - (4q, Chr11) |
| P8 | ++EGFR, +[5p15.3-p12, 5q31.1-q35.3(clone1), Chr7, 8q24], -[ 6q21-q27, Chr10, 13q13.3-q34, 15q21.2-q23(clone1), 18q21.2-q22.4], --CDKN2A/B | clone 1 selection | |
| P13 | +(Chr7, Chr19, Chr20), -(6q16.3-q21, Chr10, 17q11-12), --CDKN2A/B | - (1p21.1-p31.2, Chr13) | +(1q, 2p, 3q, 16q, Chr17), 0 (Chr10, Chr13) |
| T16 | ++[EGFR, MDM2(clone2)], +7q, - [6q, Chr10, 12q21.1-q24.3, 13q11-q31.1], --CDKN2A/B | **G1**: selection of additional clone: ++MDM4, 0 [MDM2, 12q21.1-q24.3] | 0 EGFR |
| T101 | ++[EGFR, MDM2] +[Chr1, Chr7, Chr9, Chr13, Chr17, Chr19] -[3q, Chr4, Chr10, Chr11, 14q11, Chr15], --CDKN2A/B, | **G6**: +Chr16, -Chr6 | |
| T158 | ++EGFR, +[ Chr7, Chr19, Chr20], -Chr10, --CDKN2A/B | **G1:** selection of additional clone : ++MDM2, +[12q15-q21.1], - [10p15.3-p12.31, 10q11.21-q26.3, 15q13.1-q21.3] --CDKN2A/B, 0 EGFR | - (12q24.31-q24.33), 0 (Chr19) |
| T185 | ++EGFR, +[Chr7, Chr20] –[10q, Chr22] --CDKN2A/B | | |
| T186 | complex genome ++[MYCN, CDK4] | | |
| T188 | ++EGFR, +[Chr7, Chr19, Chr20] –[1p36.31-p36.13, 6p21.32, Chr10, Chr13] --CDKN2A/B | 0 (Chr13) | |
| T192 | ++[EGFR, 2q34], +[1q21.2-24.2, Chr7, Chr19, Chr20], - [9p24.3 -13.3, Chr10] | | |
| T226 | low tumor content | ++EGFR, +(partial Chr5, Chr7), -(partial Chr1, partial Chr6, Chr10, Chr14, partial Chr16, partial Chr19, Chr22), --CDKN2A/B | |
| T233 | ++[EGFR, 2q34], +[1q21.2-24.2, Chr7, Chr19, Chr20], - [9p24.3 -13.3, Chr10] | | |
| T238 | +[Chr7, Chr19, Chr20] -[6q, Chr10, Chr13], --CDKN2A/B | | |
| T239 | +[Chr7, Chr19], -[1p36.32-p34.3, Chr10], --CDKN2A/B | | |
| T251 | ++[EGFR, 2q34], +[1q21.2-24.2, Chr7, Chr19, Chr20], - [9p24.3 -13.3, Chr10] | | |
| T281 | -CDKN2A/B; low tumor content | +Chr7, - (partial Chr1, 3p, Chr6, Chr9, Chr10, Chr13, Chr14, Chr16, partial Chr17, -CDKN2A/B) | |
| T304 | ++[EGFR, CDK4, MDM2], +[Chr1, Chr7, Chr19, Chr20], -(partial Chr1, Chr10, partial Chr12) | 0 (Chr19) | |
| T331 | ++EGFR, +[Chr1, Chr7, Chr20],- [9p, Chr10, Chr22], --CDKN2A | | |
| T341 | ++[PDGFRA, CDK6], - [3q25.3-q26.2, Chr10 ], --CDKN2A/B | | |
| T347 | ++EGFR, +[Chr7, 19p, Chr20], - [1p36.3-p34.3, 1q32.3-q44, Chr10, 13q12.11-31.1, 14q11.2-q21.3], --CDKN2A/B | | |
| T356 | low tumor content | +Chr7, -(partial Chr1, partial Chr2, partial Chr3, Chr4, Chr6, Chr10, Chr14, Ch17, Chr18) | |
| T361 | ++EGFR, + [Chr7, 19p, 20p13-p11.1, 20q13.1-q13.3], - [3q13.33-q21.1, 6p25.3-p21.32, 10q23.2-q26.3], --CDKN2A/B | | |
| T367 | ++EGFR, +[Chr7, Chr19], - Chr10 | ++MDM4 | |
| T384 | ++EGFR, low tumor content | Not assessed | Not assessed |
| T386 | + Chr7, - [1p36.3-p34.3, Chr10, 11q12.3-q13.2, 11q24.1-q13.2, 14q11.2-q24.1, 17q11.2-q21.2, 18p11.32-p11.22, Chr22 ], --CDKN2A/B | | |
| T394 | complex genome ++[PDGFRA,KIT, KDR, MET], --CDKN2A/B | | |
| T407 | complex genome ++[PDGFRA,KIT, KDR, MET], --CDKN2A/B | | |
| T434 | ++[EGFR, CDK4, MDM2], +[7p,7q36.1-q36.2, Chr20], -[1p36.23-p36.21, 6q26-q27, Chr10, 14q13-qter] | | |
| T470 | ++EGFR, +[Chr7, 19p, Chr20], - [1p36.3-p34.3, 1q32.3-q44, Chr10, 13q12.11-31.1, 14q11.2-q21.3], --CDKN2A/B | -1p31.1-p11.2 | |
| T476 | ++[EGFR, MDM4] +[1p35.2-p35.1, 1q32.1, 3q26.1-q29, Chr7], - Chr10, -- CDKN2A/B | | |
| T591 | ++EGFR, +Chr7, - [8q, partial Chr9, Chr10, 11q, Chr13, Chr15] | Not assessed | |
| T756 | complex genome | Not assessed | |
| T772 | ++EGFR, +Chr7, - Chr10, --CDKN2A/B | Not assessed | |
| T784 | +[Chr7, partial Chr3] - [partial Chr10, partial Chr13] --CDKN2A/B, additional partial chromosomal losses | Not assessed | |
| T797 | ++[PDGFRA, MDM2, CDK4], +Chr7, - [Chr4, Chr10, Chr11, partial Chr12, Chr13] | Not assessed | |
| T831/T832 | ++EGFR, +Chr7, - Chr10, --CDKN2A/B, additional partial chromosomal losses | Not assessed | |

**Table S3.** **List of genetic variants private to patient tumors and respective PDOXs.**

| | Chromosome | Region | Reference | Allele | Read count | Read coverage | Frequency | Amino acid change | Coding region change | hg19_Genes | dbSNP | AC gnomad | AF gnomad |
|---|---|---|---|---|---|---|---|---|---|---|---|---|---|
| T101_Patient | 3 | 12626516 | G | A | 56 | 250 | 22,4 | | NM_002880.3:c.1669-36C>T | RAF1 | 3729931 | 88513 | 0,35 |
| T158_Patient | 5 | 1293879 | C | A | 8 | 32 | 25 | | NM_198253.2:c.1122G>T | TERT | | | |
| T158_Patient | 9 | 139414012 | T | G | 31 | 59 | 54,1 | NP_060087.3:p.Thr250Pro | NM_017617.4:c.748A>C | NOTCH1 | | | |
| T158_Patient | 9 | 139414015 | A | C | 28 | 57 | 49,12 | NP_060087.3:p.Phe249Val | NM_017617.4:c.745T>G | NOTCH1 | | | |
| T158_Patient | 10 | 104268859..104268860 | GC | A | 49 | 137 | 37,14 | | NM_016169.3:c.183-67_183-66delinsA | SUFU | | | |
| T158_Patient | 10 | 104386934 | T | C | 50 | 113 | 44,25 | | NM_016169.3:c.1299T>C | SUFU | 17114803 | 40291 | 0,16 |
| T158_Patient | 10 | 104387019 | T | C | 45 | 114 | 39,47 | | NM_016169.3:c.1365+19T>C | SUFU | 12414407 | 165628 | 0,66 |
| T158_Patient | 10 | 123239112 | G | A | 61 | 180 | 33,89 | | NM_022970.3:c.*259C>T | FGFR2 | 1047057 | 85941 | 0,55 |
| T158_Patient | 10 | 123243197 | G | A | 45 | 106 | 42,45 | | NM_022970.3:c.2304+15C>T | FGFR2 | 2278202 | 138507 | 0,56 |
| T158_Patient | 10 | 123277507 | G | A | 51 | 171 | 29,82 | | NM_022970.3:c.1087+689C>T | FGFR2 | 117480075 | | |
| T158_Patient | 10 | 123298158 | T | C | 41 | 159 | 25,79 | | NM_022970.3:c.696A>G | FGFR2 | 1047100 | 196571 | 0,78 |
| T158_PDOX | 2 | 29451799 | T | C | 4 | 10 | 40 | | NM_004304.4:c.2766A>G | ALK | 778543123 | 7339 | 0,04 |
| T158_PDOX | 2 | 29451802 | G | C | 5 | 10 | 50 | NP_004295.2:p.Phe921Leu | NM_004304.4:c.2763C>G | ALK | 201042802 | 4651 | 0,03 |
| T158_PDOX | 7 | 142640111 | C | T | 17 | 45 | 37,78 | NP_000411.1:p.Ala598Thr | NM_000420.2:c.1792G>A | KEL | 149066842 | 32 | 1,27E-04 |
| T185_Patient | 10 | 131565064 | A | G | 20 | 71 | 28,17 | NP_002403.2:p.Ile174Val | NM_002412.4:c.520A>G | MGMT | 2308321 | 22954 | 0,09 |
| T186_Patient | 6 | 33290602 | G | C | 6 | 25 | 24 | | NM_001141969.1:c.-53+37C>G | DAXX | 1331292456 | 3448 | 0,02 |
| T192_Patient | 10 | 43622217 | T | C | 37 | 149 | 24,83 | | NM_020975.4:c.3187+47T>C | RET | 2075912 | 185804 | 0,79 |
| T233_Patient | 10 | 43595968 | A | G | 28 | 110 | 25,45 | | NM_020975.4:c.135A>G | RET | 1800858 | 1,83E+05 | 0,73 |
| T233_Patient | 10 | 43613843 | G | T | 23 | 103 | 22,33 | | NM_020975.4:c.2307G>T | RET | 1800861 | 185430 | 0,74 |
| T233_Patient | 10 | 123239112 | G | A | 35 | 111 | 31,53 | | NM_022970.3:c.*259C>T | FGFR2 | 1047057 | 85941 | 0,55 |
| T233_PDOX | 7 | 100676512 | T | G | 79 | 322 | 24,53 | NP_001035194.1:p.Ser605Arg | NM_001040105.1:c.1815T>G | MUC17 | | | |
| T233_PDOX | 7 | 148516122 | T | C | 47 | 109 | 43,12 | | NM_004456.4:c.999+566A>G | EZH2 | | | |
| T238_Patient | 6 | 87966524 | T | C | 33 | 94 | 35,11 | | NM_015021.1:c.3177T>C | ZNF292 | 143504993 | 998 | 4,02E-03 |
| T238_Patient | 6 | 114265587 | T | C | 48 | 134 | 35,82 | | NM_001527.3:c.1092-13A>G | HDAC2 | 13204445 | 57597 | 0,24 |
| T238_Patient | 9 | 2170607 | G | A | 14 | 47 | 29,79 | | NM_003070.4:c.4253+135G>A | SMARCA2 | 117087869 | | |
| T238_Patient | 9 | 5081780 | G | A | 54 | 141 | 38,3 | | NM_001322194.1:c.2490G>A | JAK2, AL161450.1 | 2230724 | 132377 | 0,53 |
| T238_Patient | 10 | 70332580 | A | G | 51 | 141 | 36,17 | NP_085128.2:p.Asp162Gly | NM_030625.2:c.485A>G | TET1 | 10823229 | 82847 | 0,33 |
| T238_Patient | 10 | 89653686 | A | G | 12 | 25 | 48 | | NM_001304718.1:c.-626-96A>G | PTEN | 1903858 | | |
| T238_Patient | 10 | 104268877 | G | C | 106 | 220 | 48,18 | | NM_016169.3:c.183-49G>C | SUFU | 2281879 | 85332 | 0,34 |
| T238_Patient | 10 | 123239112 | G | A | 99 | 308 | 32,25 | | NM_022970.3:c.*259C>T | FGFR2 | 1047057 | 85941 | 0,55 |
| T238_Patient | 10 | 123243197 | G | A | 31 | 104 | 29,81 | | NM_022970.3:c.2304+15C>T | FGFR2 | 2278202 | 138507 | 0,56 |
| T238_Patient | 10 | 123298158 | T | C | 55 | 157 | 35,03 | | NM_022970.3:c.696A>G | FGFR2 | 1047100 | 196571 | 0,78 |
| T238_Patient | 13 | 28589495 | C | G | 46 | 117 | 39,32 | | NM_004119.2:c.2654-102G>C | FLT3 | 9579142 | | |
| T238_Patient | 13 | 32890572 | G | A | 68 | 170 | 40 | | NM_000059.3:c.-26G>A | BRCA2 | 1799943 | 61394 | 0,25 |
| T238_Patient | 13 | 32911888 | A | G | 64 | 170 | 37,65 | | NM_000059.3:c.3396A>G | BRCA2 | 1801406 | 73766 | 0,29 |
| T238_Patient | 13 | 32929232 | A | G | 40 | 113 | 35,71 | | NM_000059.3:c.7242A>G | BRCA2 | 1799955 | 56617 | 0,23 |
| T238_Patient | 13 | 32936646 | T | C | 43 | 103 | 41,75 | | NM_000059.3:c.7806-14T>C | BRCA2 | 9534262 | 130837 | 0,52 |
| T238_Patient | 13 | 32953388 | T | C | 30 | 71 | 42,25 | | NM_000059.3:c.8755-66T>C | BRCA2 | 4942486 | | |
| T238_Patient | 17 | 29555971 | A | G | 28 | 74 | 37,84 | | NM_001042492.2:c.2410-72A>G | NF1 | 777315699 | | |
| T251_Patient | X | 39922282..39922285 | GAAA | - | 20 | 79 | 25,32 | NP_001116857.1:p.Leu1296fs | NM_001123385.1:c.3887_3890del | | | | |
| T331_PDOX | 7 | 100694854 | G | A | 9 | 39 | 23,08 | | NM_001040105.1:c.12875-40G>A | MUC17 | 67328257 | 74616 | 3,00E-01 |
| T331_PDOX | 9 | 5050706 | C | T | 34 | 121 | 28,1 | | NM_001322194.1:c.489C>T | JAK2 | 2230722 | 80745 | 0,32 |
| T331_PDOX | 9 | 5081780 | G | A | 33 | 97 | 34,02 | | NM_001322194.1:c.2490G>A | JAK2 | 2230724 | 132377 | 0,53 |
| T331_PDOX | 10 | 43606687 | A | G | 58 | 185 | 31,35 | | NM_020975.4:c.1296A>G | RET | 1800860 | 174813 | 0,7 |
| T331_PDOX | 22 | 21344884 | C | T | 14 | 63 | 22,22 | | NM_006767.3:c.791+70C>T | LZTR1 | 2073989 | | |
| T331_PDOX | 22 | 24167513 | G | A | 52 | 152 | 34,21 | | NM_001317946.1:c.924G>A | SMARCB1 | 2229354 | 28247 | 0,11 |
| T341_Patient | 7 | 151935962 | A | C | 9 | 30 | 30 | | NM_170606.2:c.2533-51T>G | KMT2C | 113178923 | | |
| T341_PDOX | 4 | 55138664 | G | C | 357 | 464 | 76,89 | NP_001334579.1:p.Trp460Cys | NM_001347830.1:c.1380G>C | PDGFRA | | | |
| T407_Patient | 14 | 95591070 | G | A | 21 | 57 | 36,84 | | NM_030621.4:c.904-65C>T | DICER1 | 67737119 | | |
| T407_Patient | 14 | 105239146 | C | G | 12 | 35 | 34,29 | | NM_005163.2:c.1172+69G>C | AKT1 | 3803304 | | |
| T407_Patient | 16 | 2105400 | C | T | 102 | 203 | 50,25 | | NM_000548.4:c.482-3C>T | TSC2 | 1800720 | 21836 | 0,09 |
| T407_Patient | 16 | 2115481 | C | T | 84 | 157 | 53,5 | | NM_000548.4:c.1600-39C>T | TSC2 | 45477195 | 18357 | 0,07 |
| T434_PDOX | 12 | 18762483 | G | C | 17 | 65 | 26,15 | NP_001275701.1:p.Asp1368His | NM_001288772.1:c.4102G>C | PIK3C2G | | | |
| T470_PDOX | 8 | 90948273 | C | T | 7 | 28 | 25 | | NM_001024688.2:c.1989-433G>A | NBN | 2735384 | | |
| T476_Patient | 9 | 2116037 | G | A | 12 | 48 | 25 | | NM_003070.4:c.3672G>A | SMARCA2 | 6601 | 30639 | 0,12 |

**Table S4.** **Glioma specific mutations in patient tumors and preclinical models.**
Mutations were established form targeted DNA sequencing and are displayed for a selection of glioma-relevant driver genes.
* P8 and T16* patient tumors sequenced correspond to clones 2 from aCGH

| Gene | PDOX model | P3 | P8 * | P13 | T16 * | T101 | T158 | T185 | T186 | T188 | T192 | T233 | T238 | T251 | T331 | T341 | T347 | T361 | T394 | T407 | T434 | T470 | T476 |
|---|---|---|---|---|---|---|---|---|---|---|---|---|---|---|---|---|---|---|---|---|---|---|---|
| IDH1 | Patient | | | | | | | | R132H, NM_005896 | | | | | | | | | | R132H, NM_005896 | R132H, NM_005896 | | | |
| | PDOX | | | | | | | | R132H, NM_005896 | | | | | | | | | | R132H, NM_005896 | R132H, NM_005896 | | | |
| | Cell line | | | | | | | | | | | | | | | | | | R132H, NM_005896 | R132H, NM_005896 | | | |
| IDH2 | Patient | | | | | | | | | | | | | | | | | | | | | | |
| | PDOX | | | W244R, NM_002168 | | | | | | | | | | | | | | | | | | | |
| | Cell line | | | W244R, NM_002168 | | | | | | | | | | | | | | | | | | | |
| ATRX | Patient | | S1348T, NM_000489 | | | | | | Glu1511*, NM_000489 | | | | | | | | | | D1212del E886fs, NM_000489 | D1212del E886fs, NM_000489 | | | |
| | PDOX | | S1348T, NM_000489 | | | | | | Glu1511*, NM_000489 | | | | | | | | | | D1212del E886fs, NM_000489 | D1212del E886fs, NM_000489 | | | |
| | Cell line | | | | | | | | | | | | | | | | | | D1212del E886fs, NM_000489 | D1212del E886fs, NM_000489 | | | |
| TP53 | Patient | | | | | | | | | | | | | | | | | | E204fs, NM_001126115 | E204fs, NM_001126115 | | | |
| | PDOX | C176Y, NM_001126112.2 | | | | | | | | | | | | | | | | | E204fs, NM_001126115 | E204fs, NM_001126115 | | | |
| | Cell line | C176Y, NM_001126112.2 | | | | | | | | | | | | | | | | | E204fs, NM_001126115 | E204fs, NM_001126115 | | | |
| EGFR | Patient | | A289T, NM_005228, ex10-11del | | ex2-7del | | | G598V, NM_005228 | | ex-6-7del, ex25-26del | complex rearrangements (ex1-6), ex14-15 | vIII, complex rearrangements | | vIII, complex rearrangements | R108K, NM_005228 | | ex2-15del | R108K, NM_005228 | | | F254I, NM_005228 | ex2-15del | |
| | PDOX | | A289T, NM_005228 | | ex25-27del | | | G598V, NM_005228 | | ex-6-7del, ex25-26del | complex rearrangements (ex1-6), ex14-16 | vIII, complex rearrangements | | vIII, complex rearrangements | R108K, NM_005228 | | ex2-15del | R108K, NM_005228 | | | F254I, NM_005228 | ex2-15del | G598V,NM_005228, complex rearrangements ex5-28 |
| | Cell line | | | ex23-25del | | | | | | | | | | | | | | | F254I, NM_005228 | | | | |
| MDM4 | Patient | | | | | | | | | K374Q, NM_002393 | | | T222R, NM_002393 | | K374Q, NM_002393 | | | | | | | | |
| | PDOX | | | | | | | | | K374Q, NM_002393 | | | T222R, NM_002393 | | K374Q, NM_002393 | | | | | | | | |
| | Cell line | | | | | | | | | | | | | | | | | | | | | | |
| MET | Patient | | A954S, NM_001127500 | | | | | | | | | | | | | | | | | | | | |
| | PDOX | | A954S, NM_001127500 | | | | | | | | | | | | | | | | | | | | |
| | Cell line | | | | | | | | | | | | | | | | | | | | | | |
| NF1 | Patient | | | | | | | E244G, NM_001042492 | | | | | | | | | | | | | | | |
| | PDOX | | | | | | | E244G, NM_001042492 | | | | | | | | | | | | | | | |
| | Cell line | | | | | | | | | | | | | | | | | | | | | | |
| PIK3CA | Patient | | | | | | | | | | | | | | | G118N, NM_006218 | | | | | | | |
| | PDOX | | | | | | | | | | | | | | | G118N, NM_006218 | | | | | | | |
| | Cell line | | | A990V, NM_006218 | | | | | | | | | | | | | | | | | | | |
| PIK3R1 | Patient | | | | | | G376R, NM_181523 | | | | | | | | | | | | | | | | |
| | PDOX | | | | | | | | | | | | | | | | | | | | | | |
| | Cell line | | | | | | | | | | | | | | | | | | | | | | |
| PTCH1 | Patient | | | | | | | | | V866I, NM_001083602 | | | Y1250C, NM_001083602 | | | | | | | | | | |
| | PDOX | | | | | | | | | V866I, NM_001083602 | | | Y1250C, NM_001083602 | | | | | | | | | | |
| | Cell line | | | | | | | | | | | | | | | | | | | | | | |
| PTEN | Patient | | T319fs, NM_000314 | | | | | | | | | | | | | | C304fs, NM_000314 | | | | C136Y, NM_000314 | C304fs, NM_000314 | |
| | PDOX | D236fs, NM_000314 | T319fs, NM_000314 | D107V, NM_000314 | | | | | | | | | R335*, NM_000314 | | | | C304fs, NM_000314 | | | | C136Y, NM_000314 | C304fs, NM_000314 | |
| | Cell line | L220fs, NM_000314 | | D107V, NM_000314 | | | | | | | | | R335*, NM_000314 | | | | | | | | | | |
| RET | Patient | | | | | | | | L56M, NM_020975 | | | | | | | | | | | | | | |
| | PDOX | | | | | | | | L56M, NM_020975 | | | | | | | | | | | | | | |
| | Cell line | | | | | | | | | | | | | | | | | | | | | | |
| STAG2 | Patient | | | | | | | T1151A, NM_001042749 | | | | | | | | | | | | | | | |
| | PDOX | | | | | | | T1151A, NM_001042749 | | | | | | | | | | | | | | | |
| | Cell line | | | | | | | | | | | | | | | | | | | | | | |

**Table S5.** **Classification of Glioma patient tumors and preclinical models based on the DNA methylation profiles.**

Detailed overview of the comparison of patient-, PDOX- and GSC line-related methylation-based molecular subgrouping and MGMT promoter methylation status (methylated vs. unmethylated).

Molecular diagnosis and MGMT promoter status are based on the analyses from the Heidelberg neuropathology tool https://www.molecularneuropathology.org/mnp, methylation class and methylation cluster are based on deSouza et al., 2019.

| PDOX model | Molecular diagnosis Patient | Molecular diagnosis PDOX | Molecular diagnosis cell line | MGMT methylation Patient | MGMT methylation PDOX | MGMT methylation cell line | Methylation cluster patient | Methylation cluster PDOX | Methylation cluster cell line | Methylation class patient | Methylation class PDOX | Methylation class cell line |
|---|---|---|---|---|---|---|---|---|---|---|---|---|
| P8 (clone 2*) | GBM, IDHwt, RTK I | GBM, IDHwt, RTK I/II | | unmethylated | methylated | | LGm5 | LGm4/5 | | Mesenchymal-like | Classic-like | |
| P3 | NA | GBM, IDHwt, RTK II/I | No match | methylated | methylated | methylated | NA | LGm4 | LGm4 | NA | Classic-like | Classic-like |
| P13 | NA | GBM, IDHwt, RTK II | No match | methylated | methylated | methylated | NA | LGm4 | LGm4 | NA | Classic-like | Classic-like |
| T16 (clone 2*) | GBM, IDHwt, mesenchymal | GBM, IDHwt, RTK II | No match | unmethylated | unmethylated | unmethylated | LGm5 | LGm4 | LGm4 | Mesenchymal-like | Classic-like | Classic-like |
| T101 | NA | GBM, IDHwt, RTK II | | methylated | methylated | | NA | LGm4 | | NA | Classic-like | |
| T158 | GBM, IDHwt, mesenchymal | GBM, IDHwt, RTK II | NA | unmethylated | unmethylated | NA | LGm5 | LGm4 | NA | Mesenchymal-like | Classic-like | NA |
| T185 | GBM, IDHwt, RTK II | GBM, IDHwt, RTK II | | methylated | methylated | | LGm4 | LGm4 | | Classic-like | Classic-like | |
| T186 | Glioma, IDHm, high grade astrocytoma | Glioma, IDHm, high grade astrocytoma | | methylated | methylated | | LGm1 | LGm1 | | K1: G-CIMP-low | K1: G-CIMP-low | |
| T188 | GBM, IDHwt, RTK II | GBM, IDHwt, RTK II | | unmethylated | unmethylated | | LGm4/5 | LGm4/5 | | Classic-like | Classic-like | |
| T192 | GBM, IDHwt, mesenchymal | GBM, IDHwt, RTK II | | unmethylated | unmethylated | | LGm5 | LGm5 | | Mesenchymal-like | Mesenchymal-like | |
| T226 | NA, low tumor content | GBM, IDHwt, RTK I | | NA | methylated | | | | | | | |
| T233 | GBM, IDHwt, mesenchymal | GBM, IDHwt, RTK II | | unmethylated | unmethylated | | LGm5 | LGm5 | | Mesenchymal-like | Mesenchymal-like | |
| T238 | GBM, IDHwt, inflamatory tissue | GBM, IDHwt, RTK II | | methylated | methylated | | LGm5 | LGm4/5 | | Mesenchymal-like | Classic-like | |
| T239 | GBM, IDHwt, mesenchymal | GBM, IDHwt, RTK II | | methylated | methylated | | LGm4 | LGm4 | | Classic-like | Classic-like | |
| T251 | GBM, IDHwt, RTK II/ mesenchymal | GBM, IDHwt, RTK II | | unmethylated | unmethylated | | LGm5 | LGm5 | | Mesenchymal-like | Mesenchymal-like | |
| T281 | GBM, IDHwt, mesenchymal/ RTK II | GBM, IDHwt, RTK I/II | | methylated | methylated | | | | | | | |
| T304 | GBM, IDHwt, RTK II | GBM, IDHwt, RTK I/II | | unmethylated | unmethylated | | | | | | | |
| T331 | GBM, IDHwt, mesenchymal | GBM, IDHwt, RTK II | | methylated | methylated | | LGm5 | LGm5 | | Mesenchymal-like | Mesenchymal-like | |
| T341 | GBM, IDHwt, RTK I | GBM, IDHwt, RTK I | | methylated | methylated | | LGm4/5 | LGm4/5 | | Classic-like | Classic-like | |
| T347 | GBM, IDHwt, RTK II | GBM, IDHwt, RTK II | | unmethylated | unmethylated | | LGm4 | LGm4 | | Classic-like | Classic-like | |
| T356 | GBM, IDHwt, mesenchymal | GBM, IDHwt, RTK II/I | | methylated (low tumor content) | unmethylated | | | | | | | |
| T361 | GBM, IDHwt, RTK II | GBM, IDHwt, RTK II | | methylated | methylated | | LGm4 | LGm4 | | Classic-like | Classic-like | |
| T367 | GBM, IDHwt, RTK II | GBM, IDHwt, RTK I/II | | methylated | methylated | | LGm4/5 | LGm4/5 | | Classic-like | Classic-like | |
| T386 | GBM, IDHwt, RTK I | GBM, IDHwt, RTK I | | unmethylated | unmethylated | | LGm4 | LGm4 | | Classic-like | Classic-like | |
| T394 | Glioma, IDHm, high grade astrocytoma | Glioma, IDHm, high grade astrocytoma | Glioma, IDHm, high grade astrocytoma | methylated | methylated | methylated | LGm1 | LGm1 | LGm1 | K1: G-CIMP-low | K1: G-CIMP-low | K1: G-CIMP-intermediate |
| T407 | Glioma, IDHm, high grade astrocytoma | Glioma, IDHm, high grade astrocytoma | Glioma, IDHm, high grade astrocytoma | methylated | methylated | methylated | LGm1 | LGm1 | LGm1 | K1: G-CIMP-low | K1: G-CIMP-low | K1: G-CIMP-intermediate |
| T434 | GBM, IDHwt, RTK II | GBM, IDHwt, RTK II | | unmethylated | unmethylated | | LGm4 | LGm4 | | Classic-like | Classic-like | |
| T470 | GBM, IDHwt, RTK II | GBM, IDHwt, RTK II | | unmethylated | unmethylated | | LGm4 | LGm4 | | Classic-like | Classic-like | |
| T476 | GBM, IDHwt, RTK II/I | GBM, IDHwt, RTK I/II | | unmethylated | unmethylated | | LGm5 | LGm5 | | Mesenchymal-like | Mesenchymal-like | |

*Clones not propagated in PDOXs

**Table S6** **Verhaak glioma subtypes classification.**

Classification was performed for patient tumors and respective PDOX models as well as cell lines in vitro and xenografts derived thereof.
Classification was based on initial Verhaak 2010 signatures (Verhaak et al., 2010) and the tumor intrinsic 2017 signatures (Wang et al., 2017).

| Classifier | Verhaak et al. 2010 | | Wang et al. 2017 | |
|---|---|---|---|---|
| | Patient | PDOX | Patient | PDOX |
| T16 | Neural | Classical | Classical | Classical |
| T101 | Mesenchymal | Classical | Classical | Classical |
| T185 | Mesenchymal | Classical | Classical | Classical |
| P3 | Mesenchymal | Classical | Classical | Classical |
| P8 | Neural | Proneural | Classical | Classical |
| P13 | Neural/Mesenchymal | Classical | Classical | Classical |
| | Cell line | Xenograft | Cell line | Xenograft |
| NCH421k | Proneural | Proneural | Proneural | Proneural |
| NCH644 | Proneural | Proneural | Proneural | Proneural |
| NCH601 | NA | NA | Mesenchymal/Classical | Classical |
| U87 | Mesenchymal | Mesenchymal | Mesenchymal | Mesenchymal |
| U251 | Mesenchymal | Mesenchymal | Mesenchymal | Mesenchymal |

**Table S7** **List of antibodies used in the study.**

*Flow cytometry test 106 cells/100µl

| Epitope | Conjugate | Species reactivity | Clone | Supplier | Concentration used/test* |
|---|---|---|---|---|---|
| CD31 | - | mouse | 390 | Millipore | IHC:1/200 |
| Nestin | - | human | 10C2 | Millipore | IHC:1/200 |
| Vimentin | - | mouse/rat/human | EPR3776 | Epitomics | IHC:1/200 |
| Ki67 | - | mouse/rat/human | SP6 | ThermoScientific | IHC: 1/100 |
| H2AX-P | - | mouse/rat/human | 9718T | Cell Signaling | IHC: 1/500 |
| Anti-mouse -IgG | HRP | mouse | | GE Healthcare LNA931V/AG | WB: 1/10 000 |
| Goat anti-rat IgG | Alexa Fluor 555 | rat | | Invitrogen | IHC:1/1000 |
| Anti-mouse IgG | HRP | mouse | | Dakocytomation | Kit concentration |
| Anti-rabbit IgG | HRP | rabbit | | Dakocytomation | Kit concentration |
| Anti-Mouse IgG | Biotinylated | horse | | Vestor labs, BA-2000 | IHC:0.2ug/ml |
| CD15/SSEA-1 | Alexa Fluor 647 | human | MC-480 | Biolegend | 5µl/test |
| CD31 | Dy590 (PE-TR) | human | MEM-05 | Immunotools | 10µl/test |
| CD44 | PE-Cy7 | human/mouse | IM7 | eBioscience | 1.2µl/test |
| CD45 | PE-Cy7 | human | HI30 | Immunotools | 5µl/test |
| CD90 | PECy7/APC | human | 5,00E+10 | BD Bioscience | 5µl/test |
| CD133 | PE /APC | human | 293C3/AC 133 | Miltenyi | 10µl/test |
| A2B5 | APC/PE | human/mouse | 105-HB29 | Miltenyi | 10µl/test |
| EGFR | PE | human | EGFR.1 | BD Bioscience | 20µl/test |
| Lamin A/C | PE | human | sc-7292 | Santa Cruz | 20µl/test |
| EGFR | - | human | ocktail R19/ | ThermoScientific | WB: 1/1000 |
| GAPDH | - | human | D16H11 | Cell Signaling | WB: 1/1000 |
